# Isolation of phages infecting the zoonotic pathogen *Streptococcus suis* reveals novel structural and genomic characteristics

**DOI:** 10.1101/2025.01.07.631744

**Authors:** Emmanuel Kuffour Osei, Reuben O’Hea, Christian Cambillau, Ankita Athalye, Frank Hille, Charles M.A.P. Franz, Áine O’Doherty, Margaret Wilson, Gemma G. R Murray, Lucy A. Weinert, Edgar Garcia Manzanilla, Jennifer Mahony, John G Kenny

## Abstract

Bacteriophage research has experienced a renaissance in recent years, owing to their therapeutic potential and versatility in biotechnology, particularly in combating antibiotic resistant-bacteria along the farm-to-fork continuum. However, certain pathogens remain underexplored as targets for phage therapy, including the zoonotic pathogen *Streptococcus suis* which causes infections in pigs and humans. Despite global efforts, the genome of only one infective *S. suis* phage has been described. Here, we report the isolation of two phages that infect *S. suis*: Bonnie and Clyde. The phages infect 58% of 100 *S. suis* strains tested, including representatives of seven different serotypes and thirteen known sequence types from diverse geographical origins. Clyde suppressed bacterial growth *in vitro* within two multi-strain mixes designed to simulate a polyclonal *S. suis* infection. Both phages demonstrated stability across various temperatures and pH levels, highlighting their potential to withstand storage conditions and maintain viability in delivery formulations. Genome comparisons revealed that neither phage shares significant nucleotide identity with any cultivated phages in the NCBI database and thereby represent novel species belonging to two distinct novel genera. This study is the first to investigate the adhesion devices of *S. suis* infecting phages. Structure prediction and analysis of adhesion devices with AlphaFold2 revealed two distinct lineages of *S. suis* phages: *Streptococcus thermophilus*-like (Bonnie) and *S. suis*-like (Clyde). The structural similarities between the adhesion devices of Bonnie and *S. thermophilus* phages, despite the lack of nucleotide similarity and differing ecological niches, suggest a common ancestor or convergent evolution, highlighting evolutionary links between pathogenic and non-pathogenic streptococcal species. These findings provide valuable insights into the genetic and phenotypic characteristics of phages that can infect *S. suis*, providing new data for the therapeutic application of phages in a One Health context.

## Introduction

The intensification of livestock farming systems has been predicted to promote the emergence of pathogens from within the microbiota of host populations (1). One such bacterium is *Streptococcus suis*, a ubiquitous coloniser of the upper porcine respiratory tract. *S. suis* is a secondary pathogen in the porcine respiratory diseases complex, a polymicrobial syndrome that affects the respiratory system of pigs. However, *S. suis* is capable of systemic dissemination resulting in diseases such as meningitis, endocarditis, arthritis, and septicaemia (2).Furthermore, the threat posed by *S. suis* is not limited to porcine hosts. Less than 15 years after its discovery in pigs in 1954, *S. suis* was implicated in a case of human meningitis in Denmark (3). Subsequently, several outbreaks of human infections with similar clinical manifestations in pigs have been reported globally (4). Zoonotic transmission of the pathogen to humans occurs via direct contact with infected pigs, or consumption of undercooked pork products.

Although only partially understood, the pathogenicity and virulence of *S. suis* has been linked to over 100 putative virulence factors including suilysin (*sly*), muramidase-release protein (*mrp*), extracellular protein factor (*epf*), and capsular polysaccharides (*cps*) (5). CPS is a critical virulence factor involved in evasion of host immune mechanisms and is also the basis on which the bacterium is classified into 29 serotypes (6). Serotype 2 is reported as the most common cause of *S. suis* infection in pigs and humans globally. However, at the regional level, strains of serotypes ½, 9, and 1 are the most common agents of *S. suis* infections in South America, Western Europe, and North America, respectively (7). Although vaccine candidates exist, efforts to produce a universal vaccine that is cross-protective is confounded by the genetic diversity of the species (8), particularly in the *cps* loci, a common antigenic target for vaccine development. Additionally, the *cps* clusters are prone to recombination, which can result in capsular switching (9). This implies that virulent strains can evolve *cps* types not targeted by vaccines.

Antibiotics remain the primary control strategy against the pathogen, which has contributed to the emergence and dissemination of antibiotic-resistant strains. Previous studies that examined antimicrobial resistance (AMR) in *S. suis* reported high levels of resistance to drugs including those not used in treating *S. suis* infections (10). Furthermore, several studies have reported high carriage of AMR determinants in *S. suis* genomes (11–13). The impending disaster of AMR has led to increased interest in the potential application of bacteriophages (phages) as an alternative/adjunct to antibiotics. Phages are viruses that can infect and kill bacteria. With an estimated abundance of 10^30^ to 10^32^ particles in the biosphere, these ubiquitous biological entities have been isolated from a wide range of environments (14). As the international community moves towards a One Health approach to tackling AMR along the farm-to-fork continuum, coupled with the consumers’ growing preference for greener, antibiotic-free, sustainably farmed products, phages have emerged as a promising candidate for controlling bacterial pathogens. Phages target bacteria in a highly host-specific manner and have no reported serious adverse effects on treated subjects. As such, they have been evaluated for their potential use *in vivo* within laboratory settings, agricultural sites, and in medicine for various pathogens (15–18). However, certain bacteria including *Gardnerella vaginalis*, *Clostridioides difficile*, *Porphyromonas gingivalis*, and *S. suis* remain poorly explored as targets for phage therapy, often due to difficulties in isolating virulent phages against them. Despite global efforts, to date, only one “infective” phage (phage SMP) against *S. suis* has been isolated and genome-sequenced (19) while temperate phages have also been induced from *S. suis* strains (20). Despite the setbacks in isolating and characterising virulent phages, (pro)phage-derived lysins have been developed and evaluated for bactericidal activity against *S. suis*, as with *Gardnerella vaginalis* and *Clostridium difficile* (21–24). Thus, it is imperative to characterise temperate phages and explore their potential use in bacterial control using their derivatives such as phage-derived lysins or engineered phages. Furthermore, it is important to study temperate phages as phage resistance mechanisms have been linked to *S. suis* virulence (25).

In this study, we describe the isolation of two phages named Bonnie and Clyde, tripling the number of previously characterised phages infecting *S. suis*. The genomes of both phages were sequenced and compared with phage SMP and other close relatives, revealing both phages belong to distinct, novel genera. We also describe the phage structures and adhesion devices using electron microscopy and *in silico* predictions, respectively. Furthermore, we demonstrate that the two phages can infect and lyse *S. suis* strains of various serotypes and sequence types (STs) isolated from different countries.

## Materials and methods

### Bacterial strains

All *Streptococcus suis* strains used in this study are listed in Table S1. *S. suis* 21171_DNR38 and 19867_M106485_R39 are serotype 2 strains isolated from the respiratory tract of a diseased pigs in Denmark and Spain, respectively (1). All strains and phages were cultivated in Todd Hewitt broth (THB) (Neogen, #NCM0061B) or agar (1.5% w/v) at 37°C under microaerophilic conditions. Serotypes of strains were determined using a two-step multiplex PCR (Table S2) (26) or *in silico* prediction based on whole genome sequencing (27). Multilocus sequence typing was performed with PubMLST (28) and new allele sequences and unassigned MLST profiles were submitted to PubMLST for assignment. In total, 26 novel STs have been identified and subsequently added to the *S. suis* PubMLST database.

### *S. suis* phage screening and isolation

Thirty samples including oral fluids from healthy pigs across 20 farms in Ireland, along with post-mortem lung tissue from diseased pigs, were collected and processed. Oral fluids were supplemented with 0.5 M sodium chloride (NaCl) and centrifuged at 5,000 ×g for 15 minutes. Up to 10 g of lung tissue was homogenised in a stomacher for 10 minutes in 90 mL of sterile SM buffer (50 mM Tris-HCl, 0.1 M NaCl, and 8 mM MgSO₄ [pH 7.4]) and centrifuged at 5,000 ×g for 15 minutes. The supernatants from the oral fluids and tissue homogenates were filtered through a 0.45 μm polyethersulfone membrane filter (Sarstedt), and the resulting filtrates were screened for phages using the double-layer agar (DLA) plaque assay method (1.5% agar [w/v] underlay and 0.8% agar [w/v] overlay) (29). A total of 50 strains were used in screening, with serotypes 2, 9, and 14 more highly represented due to their frequent association with infections (Table S1).

### *S. suis* phage propagations and purification

To obtain a homogeneous phage lysate, five rounds of single plaque purification were performed, all carried out on strain 21171_DNR38. Subsequently, to generate phage lysates, a clonal plaque was used for plaque assays to generate confluent lysis. Then SM buffer was added to twelve such plates and the plates were incubated on a shaker at 100 rpm for 4 hours. The top layer was disrupted, and the resulting slurry was aspirated into a sterile tube. The lysate was collected by centrifugation at 4,500 ×g for 15 minutes and filtered through a 0.45 µm filter.

For liquid propagation, an exponentially growing culture (OD600 of 0.2) supplemented with 40 mM MgCl₂ and 1 mM CaCl₂ was infected with 0.02 volumes of the plaque-purified lysate and incubated overnight. The culture was then centrifuged, filtered, and the titre was estimated using a DLA spot assay method. Briefly, 10 µL of 10-fold serially diluted lysate was spotted on double-layer plates and incubated overnight at 37°C under aerobic conditions. The resulting plaques were counted, and the titre was expressed as PFU/mL.

### Phage DNA extraction and genome sequencing

The Norgen Biotek phage DNA isolation kit was used to extract DNA from the purified phage lysates with some modifications to the manufacturer’s instructions (Norgen Biotek Corp., Ontario, Canada). Briefly, 1 mL of the phage lysate (10^8^ PFU/mL) was treated with 20U of DNase and 10X DNase buffer (#AM2238), and 1 μL of RNaseA (#EN0531). The reaction was incubated at 37°C for 70 minutes followed by nuclease inactivation with 20mM EDTA. Four microliters of proteinase K (20 mg/mL) and 500 μL of lysis buffer was added and vortexed for 10 seconds. This was incubated in a water bath at 56°C for 30 minutes. The mixture was treated with 320 μL isopropanol (Fisher Bioreagents) and purified on a column according to the manufacturer’s instructions. Phage genome libraries were prepared using the Nextera XT Library Preparation Kit and sequenced on an Illumina MiSeq sequencing system, generating 2 x 250 paired-end reads (executed by GenProbio s.r.l, Italy).

### Prophage prediction

The genome sequence of the bacterial host strain (19867_M106485_R39, accession no. DARZKS000000000.1) was screened for the presence of prophages using geNomad with the end-to-end command (30). Nucleotide sequences corresponding to ‘provirus’ topology hits were extracted and annotated with pharokka using default parameters (31). The annotated genomes were manually inspected for hallmark genes using keywords such as “terminase”, “capsid”, “tail” and “holin”.

### Prophage induction

Prophage induction from exponentially growing cultures of 19867_M106485_R39 was performed using mitomycin C (MitC), D-L-threonine coupled with temperature cycling, or UV light exposure.

For UV induction, 38 mL of THB supplemented with 10 mM MgSO₄ was inoculated with 2 mL of overnight culture and grown to an OD600 of 0.2. Subsequently, 6.5 mL aliquots of the culture were transferred into 90 mm petri dishes, achieving a depth of approximately 1 mm. The cultures were irradiated with a germicidal UV-C lamp at 253.7 nm for either 30 seconds or 2 minutes. After irradiation, the cultures were pooled and incubated at 37°C for 2 hours. The cultures were then centrifuged at 4,000 ×g for 10 minutes at 4°C and filtered through a 0.45 μm filter. An aliquot of the lysate was taken to visualise the presence of plaques. As low levels of induction were expected, the induction lysate was also enriched by adding 50 μL of host (21171_DNR38) culture to 1 mL of lysate, followed by overnight incubation. The propagation mix was filtered and stored at 4°C.

For mitomycin C induction, MitC was added to a 10 mL exponential-phase (OD600nm of 0.2) culture at a final concentration of 1.5 μg/mL and incubated for 15 hours. The induction mix was centrifuged, filtered, and enriched as described above.

By serendipity, an unintended prophage induction was observed when a culture was moved from 37 to 4 °C incubation overnight. Consequently, we attempted inducing the same prophage using controlled rapid temperature cycling and D-L-threonine supplementation. Briefly, 9.5 mL of THB supplemented with 40 mM D-L-threonine and 10 mM MgSO₄ was inoculated with 500 μL of overnight culture and incubated at 37°C for 3 hours, moved to 56°C for 2 minutes and then incubated at 4°C overnight. The following day, the culture was removed from 4°C and immediately incubated at 37°C for 3 hours, followed by 1 hour at 4°C. The culture was then centrifuged, filtered, and enriched.

All induction lysates were screened for plaque formation on 21171_DNR38 using the DLA spot assay as previously described. Further confirmation of successful induction was determined by amplifying the gene that encodes the terminase large subunit of the predicted prophage from the 19867_M106485_R39 strain using specific PCR primers (Table 1). The induction mixture was treated with DNase and RNase to remove host and unpackaged phage nucleic acid fragments followed by inactivation of the nucleases as described above. Universal *16S* rRNA gene primer pair was used as control to detect chromosomal, non-phage bacterial gene (26). Lysates from all three methods of induction were subjected to this PCR using PCR Master Mix 2X (#K0172, ThermoFisher Scientific) according to manufacturer’s instructions (Table S3, Table 1).

**Table 1:**
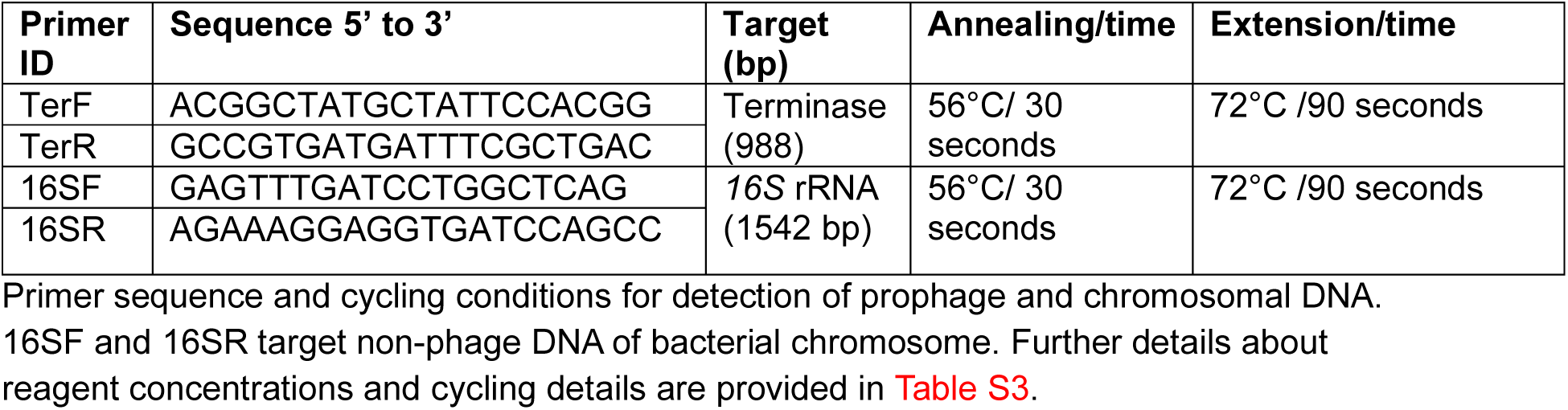
Primer sequences and PCR conditions for phage detection.

### Genome assembly and annotation

Phage genome *de novo* assembly was carried out using the SPAdes-based genome assembler Shovill (32). The --trim flag and --depth 100 flag were invoked to remove Illumina adaptors and reads were subsampled to a predicted 100x coverage. The assembled contigs were checked for completeness using CheckV1.0.1 (33). Genome annotation was performed with Pharokka v1.7.3 (31) and RAST pipeline (34). The pharokka pipeline uses PHANOTATE and Prodigal for gene prediction and assigns functional annotations using MMseqs2 by matching predicted coding sequence (CDS) to the PHROGs, VFDB and CARD databases. The --dnaapler command was used to reorient phage genomes to begin with the large terminase subunit-encoding gene. To improve annotations, the genbank (.gbk) output from pharokka was set as input for phold, a tool which utilises foldseek and colabfold to predict structural homology (35). Lifestyle of phages was predicted with PhageTYP (36). Circular genome maps of phages were constructed with phold plot command. Auxilliary metabolic genes (AMGs) encoded in phage genomes were screened using DRAM-v and VIBRANT (37, 38). An *in silico* screen of anti-viral defense systems was performed using DefenseFinder (39). Protein sequences of phage endolysins were extracted and concatenated for alignment with Clustal Omega (40). Seqvisr (41) was used to visualise amino acid similarity using SMP endolysin as reference.

### Phylogeny and comparative analysis of *S. suis* (pro)phages

Genomes of (pro)phages that share significant nucleotide homology with the phages isolated in this study were identified with BLASTN using the core nucleotide database in two searches. For both searches, only sequences with ≥50% query coverage were considered in subsequent analyses. In the first search, the taxid 1307 was excluded to filter out bacterial (i.e. *S. suis*) hits. This returned few viral (prophage) sequences that met the criteria. The search was repeated without the filter and top hits were bacterial (Table 2). The regions in the bacterial genomes (prophages) that share significant homology with the phages were extracted with geNomad as described above. A viral proteomic tree was generated in VipTree using the “with reference” setting (42). The tree was constructed based on tBLASTx computations of genome-wide similarities. Intergenomic similarities among the phages and their closest relatives were estimated using VIRIDIC. PhageGCN and taxmyPHAGE pipelines were used to predict phage taxonomic classification (36, 43). VICTOR (https://ggdc.dsmz.de/victor.php) was used to generate a phylogenomic tree. It estimated the nucleotide pairwise similarity using the Genome-BLAST Distance Phylogeny (GBDP) method. The computed intergenomic distances were used to infer a balanced minimum evolution tree with branch support via FASTME including subtree pruning and regrafting postprocessing for the D4 formula. Branch support was inferred from 100 pseudo-bootstrap replicates each. VirClust was used for orthologous genes prediction, virus clustering and the estimation of core proteins (44).

**Table 2:**
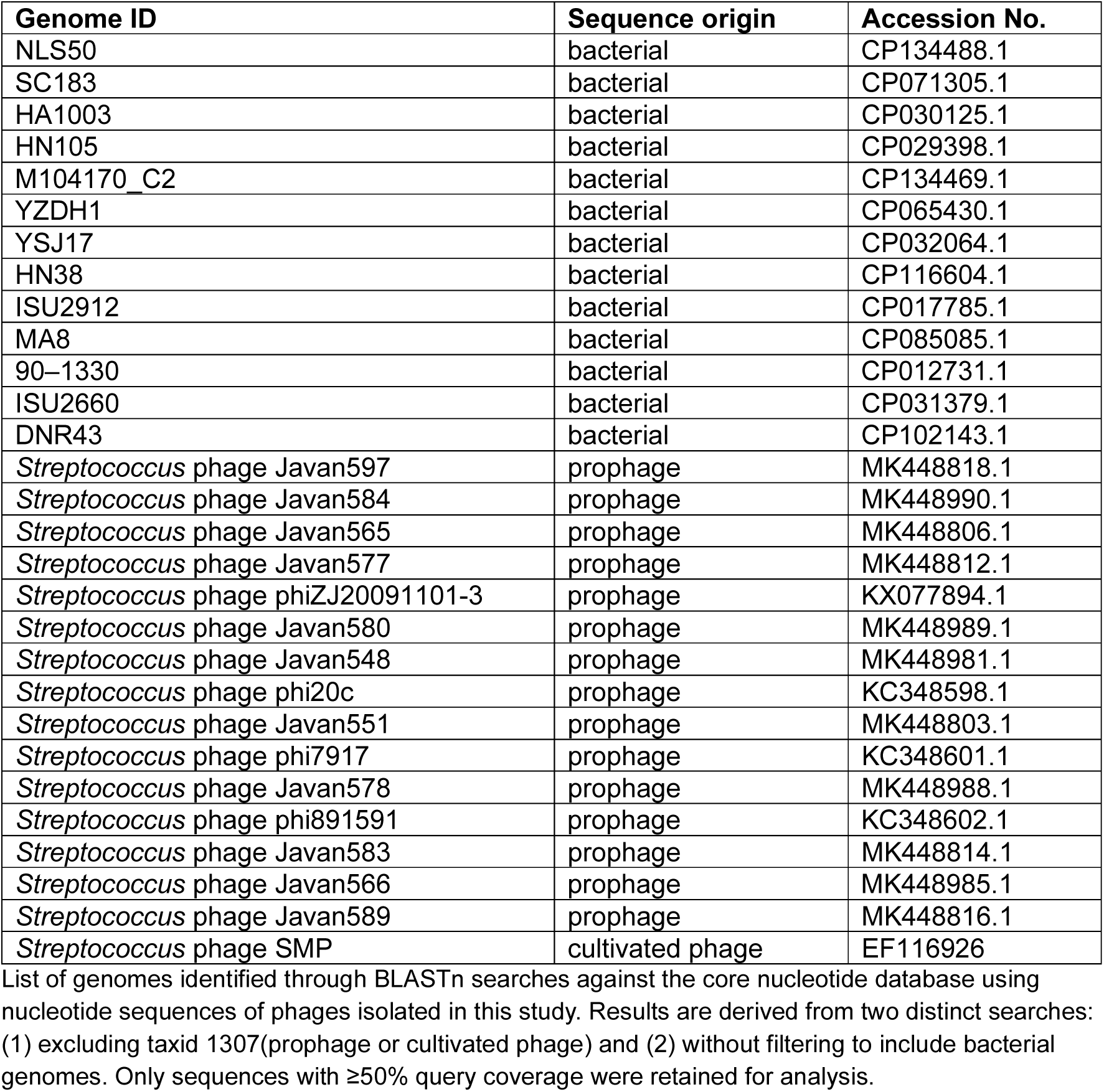
Summary of sources of (pro)phage sequences used in phylogenomic and comparative analyses.

### Host range analysis, efficiency of plating, and phage killing assay

To determine the host range of the two *S. suis* phages, 10 µL of lysate was spotted onto a double-layer agar lawn of each of the 100 host strains (Table S1). The plates were incubated overnight at 37°C. Host sensitivity was scored as complete/turbid lysis (+) or no lysis (-). Host sensitivity was confirmed by determining the efficiency of plating (EOP). Diluted phages were spotted on strains shown to be sensitive to a phage, and the plaques counted. The results were expressed as the titre of phages on the test strain relative to the titre on the original host strain.

The *in vitro* lytic activity of phages on sensitive strains was assessed by infecting exponentially growing host cells with phages at multiplicities of infection (MOI) of 0.1, 1, 10 and 100. Phages were serially diluted in a 96-well plate and equilibrated by incubating the plate at 37°C for 30 minutes. The phage dilutions were then inoculated with bacteria to a final concentration of 10^7^ CFU/mL. Prior to each reading, the plate was shaken for 5 seconds, and the absorbance at OD600nm was measured every 10 minutes over 24 hours using a microplate reader (Biotek Synergy HT Plate Reader, USA). Wells containing only phage suspension or only bacterial culture served as controls. The results are presented as the average of three independent replicates. A threshold of *p* < 0.05 was considered statistically significant.

### Thermal and pH stability

The temperature stability of *S. suis* phages was evaluated by incubating phage lysates at 4, 37, 40, 45, 50, 60, or 70°C for 1 hour, followed by estimating the titres using DLA spot assays. For pH stability, the pH of SM buffer was adjusted with either NaOH or HCl to obtain a pH of 2, 3, 4, 5, 6, 7, 8, 9, 10, 11, 12, and 13. Phage pH stability was assessed by adding 50 μL of ≥10^8^ PFU/mL phage to 450 μL of pH-adjusted SM buffer. The mixture was incubated at 37°C for 1 hour, 2 hours or 24 hours, and the titres were estimated using the DLA method.

### One-step growth curve

The latent period and burst size of the phages were determined in a one-step growth experiment as described by Kropinski (45). Briefly, a 10 mL exponential-phase host culture was centrifuged and resuspended in 900 μL of THB. Phage lysate was added at a final concentration of MOI 0.01 and allowed to adsorb for 15 minutes at 37°C. Unadsorbed phages were removed by centrifugation, and the pellet was resuspended in 10 mL prewarmed medium. The mixture was incubated at 37°C for 2 hours, with 200 μL aliquots taken every 5 minutes, centrifuged and the supernatant diluted to estimate phage titres. The calculated PFU/mL was plotted against time and burst size and latent period were estimated from the graph (45).

### Transmission electron microscopy

Two litres of phage lysate of Bonnie and Clyde was centrifuged to remove debris, filtered, treated with 10% (w/v) polyethylene glycol 8,000, and incubated at 4°C overnight. The phage particles were harvested by centrifugation (10,000 ×g, 15 min) and resuspended in 4 mL of SM buffer. An equal volume of chloroform was used to extract phages from the PEG suspension three times. Purification of phage lysate was performed by loading the phage on a discontinuous cesium chloride (CsCl) gradient (5M and 3M) followed by centrifugation at 13,170 ×g for 2.4 hours at 4°C (Beckman Coulter Optima L-90K series). Phage bands were extracted and dialysed against SM buffer (50 mM Tris-HCl, 0.1 M NaCl, and 8 mM MgSO4 [pH 7.5]). Phages were pipetted on 100-mesh copper grids coated with carbon and allowed to adsorb for 20 minutes. Subsequently, the grids were washed twice with deionised water and negatively stained with 2% uranyl acetate for approximately 20 seconds. Electron micrographs were generated on a Talos L120C transmission electron microscope using a 4 k x 4 k Ceta camera set to an acceleration voltage of 80 kV. Phage size was measured using the TEM analysis software Velox v3.9.0 (all equipment from ThermoFisher Scientific, Eindhoven, Netherlands) and the mean and standard deviation of at least 10 measured phage particles were calculated. The TEM images were manually improved using GIMP v2.10.32 for contrast and brightness adjustment.

### AlphaFold prediction and analysis of adhesion devices

Structures of the adhesion device proteins (Dit, Tal and/or RBP) of Bonnie and Clyde were predicted using AlphaFold2 as well as those of SMP using AlphaFold3 via google servers at https://golgi.sandbox.google.com (46, 47). Stretches of Tal with overlapping segments were predicted to allow the assembly of full-length multimers using Coot (48). pLDDT and PAE values are reported in Fig. S1. The final predicted domain structures were submitted to the Foldseek server to identify closest structural homologs in the PDB (49). Sequence alignments were performed with Multalin (50). Visual representation of the structures were prepared with ChimeraX (51).

### Statistical analysis

All quantitative datasets generated from *in vitro* experiments were analysed by estimating mean and standard error of the mean using independent biological replicates, each conducted on different days with different bacterial cultures and fresh phage lysates. GraphPad Prism 10.2.0 (USA) was used for the statistical analysis. Two-way analysis of variance (ANOVA) with Dunnett’s multiple comparisons test was used to evaluate temperature stability data at a statistical significance level of 0.05. The statistical significance of pH stability data was estimated using Kruskal-Wallis test. For phage *in vitro* lytic activity, one-way ANOVA with Tukey’s post-hoc test was performed to determine statistically significant differences. The comparisons were made between the test MOIs and the no-phage control.

## Results

### Isolation of *S. suis* infecting Phages, Bonnie and Clyde

Over 30 samples collected from pig farms across Ireland, including slurry, oral fluids, lung and tonsil tissues were processed and screened for phages against 50 *S. suis* strains, representing different serotypes. Initially, one phage was isolated from a post-mortem lung sample. This phage was initially plaque-purified on strain 19867_M106485_R39 and subsequently sequenced. However, sequencing results revealed the presence of two contigs in the supposed single phage lysate with contig coverages of 67.3 and 36.9 and sizes of 36,310 bp and 34,743 bp, respectively. CheckV and pharokka annotations confirmed that the contigs represented two distinct phages. To investigate the source of the two phages, we first screened the genome of the propagation host for prophages using geNomad. Two “provirus” topology hits were predicted: a 34 kb prophage and a 13 kb region (Table S4). Nucleotide BLAST (core nucleotide database) results of the 34 kb prophage region showed a 100% match to the second contig. This indicated that the 34 kb prophage was induced from the propagation host. To isolate the two phages individually, a suitable host strain that would minimise the risk of induction was needed. Accordingly, we screened the genomes of susceptible hosts to identify strains without prophages or with prophage-elements that lack hallmark genes including capsid and tail proteins. Strain 21171_DNR38 was found to harbour a 13 kb provirus that encodes no structural genes and was therefore used for further plaque purification. Following 22 successive passages on strain 21171_DNR38, clonal plaques corresponding to the first phage of 36 kb (vB_SsuS-Bonnie hereinafter referred to as Bonnie) were obtained. This was initially verified by PCR using primer pairs that targeted genes encoded by either of the two co-propagated phages (Table S3). The second phage was (re)isolated by inducing it from its host (19867_M106485_R39) through UV-light exposure, MitC induction, or temperature cycling. Lysates emanating from all three methods produced translucent plaques, however, following enrichment, clear spots and transparent plaques were observed (Fig. 1A). Furthermore, PCR was used for the preliminary identification of the phage in the induction lysate, matching it to both the predicted prophage and the second phage/contig (Fig. 1B), vB_SsuS-Clyde (hereinafter referred to as Clyde). The names were inspired by Bonnie Parker and Clyde Barrow, reflecting how inseparable the phages were on strain 19867_M106485_R39. All three methods for inducing phage Clyde were successful; however, UV induction-derived lysate was used for subsequent experiments based on the higher titres achieved after the enrichment.

**Fig. 1:**
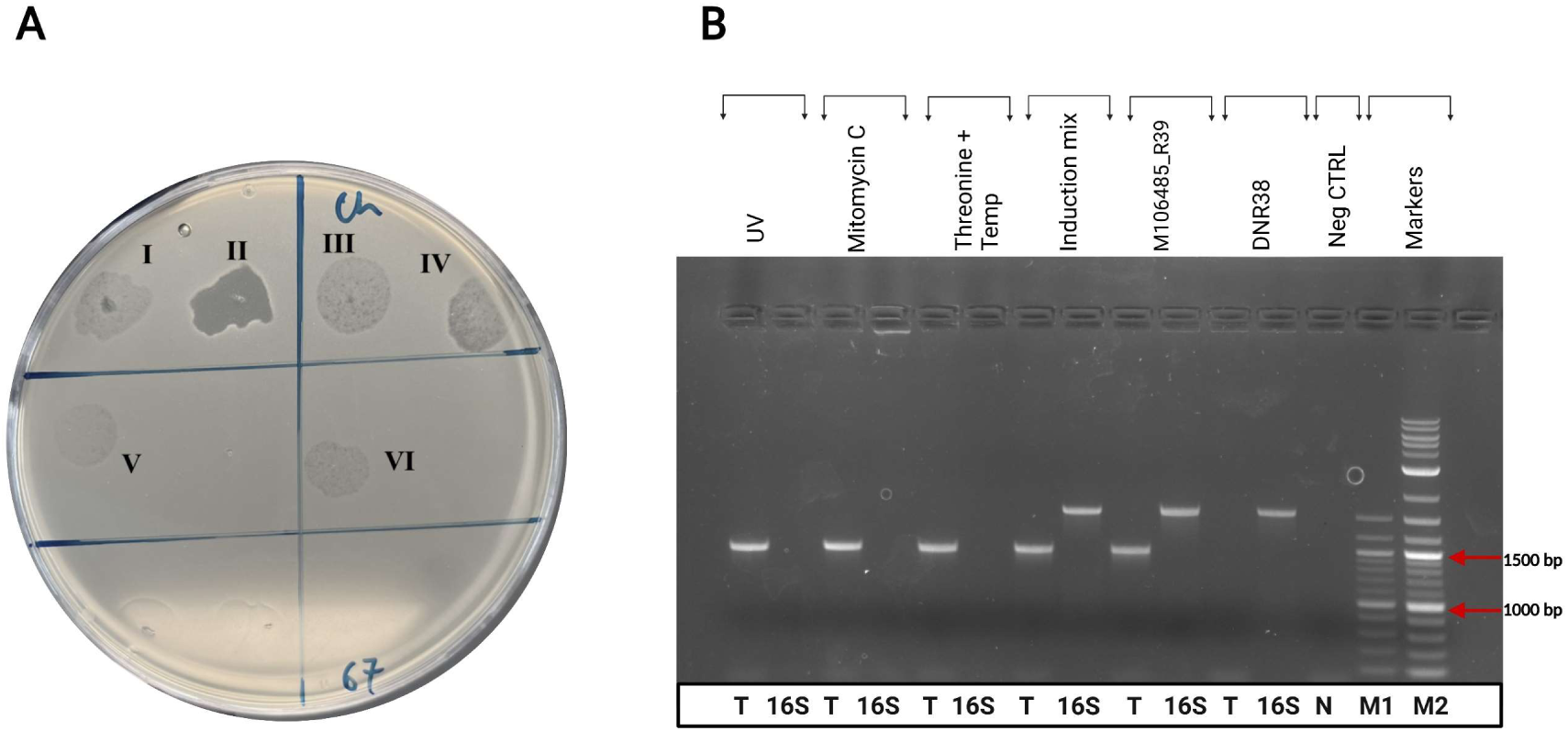
Confirmation of induction of phage Clyde by three methods **(A)** Spot assays showing filtered induction lysates on sensitive strain 21171_DNR38 before (I, III, V) and after enrichment (II, IV, VI) on overlay plates for UV, MitC, and temperature cycling prophage induction, respectively. (**B)** PCR products of terminase (T) and *16S* rRNA (16S) gene amplified from filtered, nuclease-treated lysates induced by UV, MitC, and temperature-dependent threonine induction. A crude induction mixture (non-filtered and non-nuclease treated) and a culture of host strain 19867_M106485_R39 were used as positive controls. Negative controls included strain 21171_DNR38, which does not harbour the expected prophage, and water. M1 and M2 indicate 100 bp and 1 kb DNA ladders, respectively.

### Bonnie and Clyde are temperate phages

Genomic DNA was extracted from purified lysates of Bonnie and Clyde, sequenced, and assembled into genomes of 36,160 bp and 34,734 bp, with average GC content of 40.51 and 41.87%, respectively, similar to that of the propagation host (41.44%). Bonnie has a coding density of 96.74%, encoding 72 coding sequences (CDS) of which 44 could not be assigned a function by pharokka. Clyde has coding density of 95.07%, which encodes 64 CDS with 41 of unknown function. CDS annotations improved following application of the structural homology tool, PHOLD. The number of proteins with assigned function improved from 28 to 35 and 23 to 33 for Bonnie and Clyde, respectively.

The CDSs with assigned functions could be grouped into structural (tail and capsid), lysis, integration & excision, nucleotide metabolism, and transcriptional regulation.

Neither phage genome was predicted to encode tRNAs, AMR genes, toxins, or virulence factors. However, integrases, repressors, and other genes involved in lysogeny-lysis decision making were identified in both genomes. The lifestyle of the phages was further confirmed as temperate based on PhageTYP predictions (Table S5). Moreover, DefenseFinder identified Retron_VII_2 DUF3800, one of two genes involved in Retron Type VII anti-viral system in Clyde implying a role in superinfection immunity. A DNA methyltransferase involved in the cytosine and methionine metabolism pathways was detected within the nucleotide metabolism module of Bonnie using the VIBRANT pipeline. No AMGs were identified in Clyde.

**Fig. 2:**
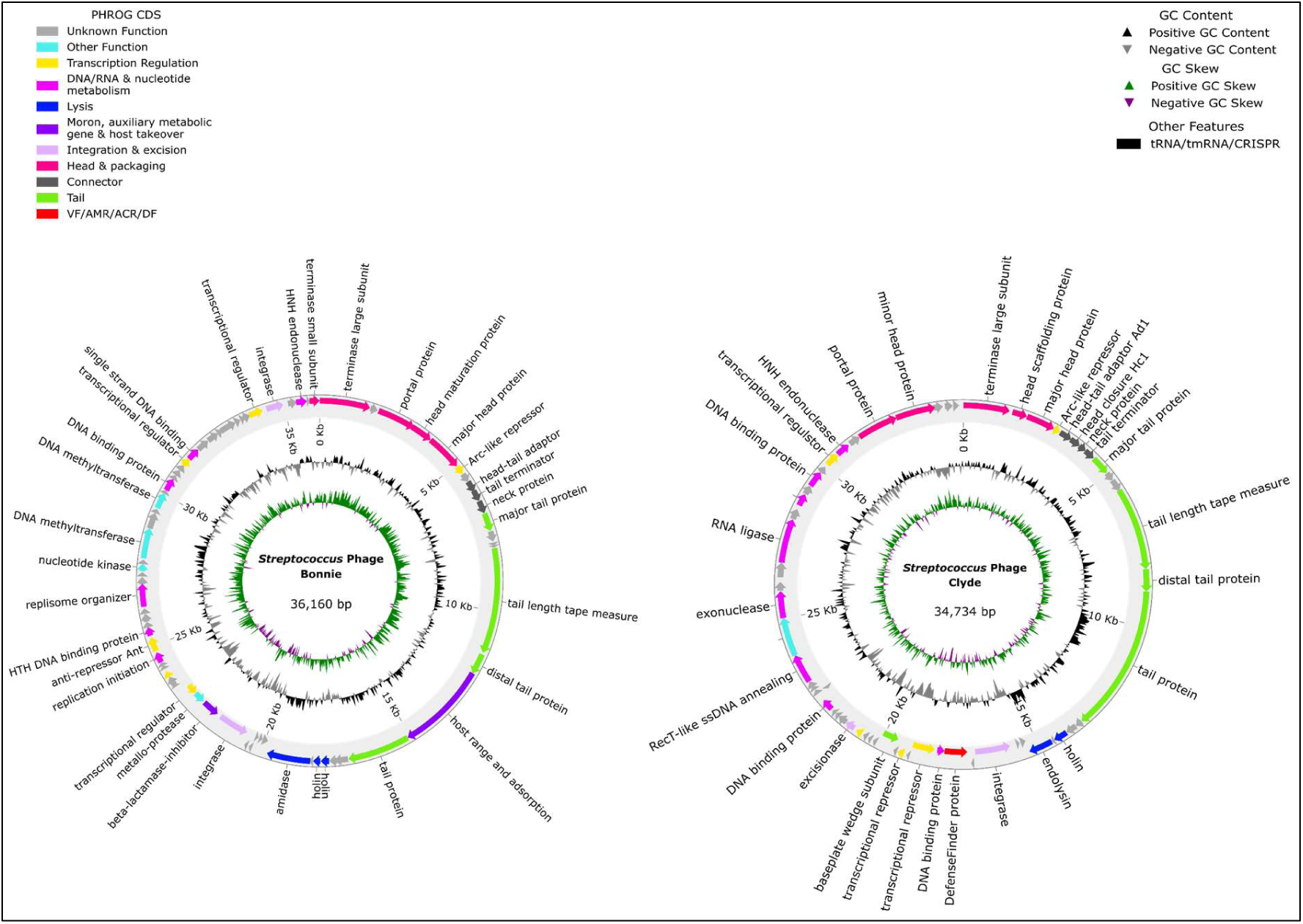
Circular genome map of *S. suis* phages generated with PHOLD. **(A)** Bonnie and **(B)** Clyde. CDSs are represented as arrows indicated in the direction they are encoded. Arrows are colour-coded based on function. Nucleotide sequences of Bonnie and Clyde are available on GenBank under accession numbers PQ0431 and PQ0432, respectively.

### Phylogeny and comparative analysis of Bonnie and Clyde

A viral proteomic tree was constructed using the nucleotide sequences of both phages in ViPtree. The resulting tree included 1,083 dsDNA viruses including Bonnie and Clyde (Fig. 3). Bonnie clustered closely with *Streptococcus* prophage phiD12 and four phages infecting *S. pneumonia* and *S. oralis*. Although distant, Clyde clustered with the *S. suis* phage SMP but more closely with *S. suis*-infecting prophage phi20c. Other members within the same clade include phages that infect *Lactococcus lactis*, *S. parauberis*, *S. mitis* and *S. pyogenes*. The genome size of all the related phages (Fig. 3) ranged from 31–49 kb. Taxmyphage and PhageGCN placed Bonnie and Clyde in the class level (Caudoviricetes). Neither phage could be classified into any genus or species recognised by the International Committee on Taxonomy of Viruses (ICTV).

**Fig. 3:**
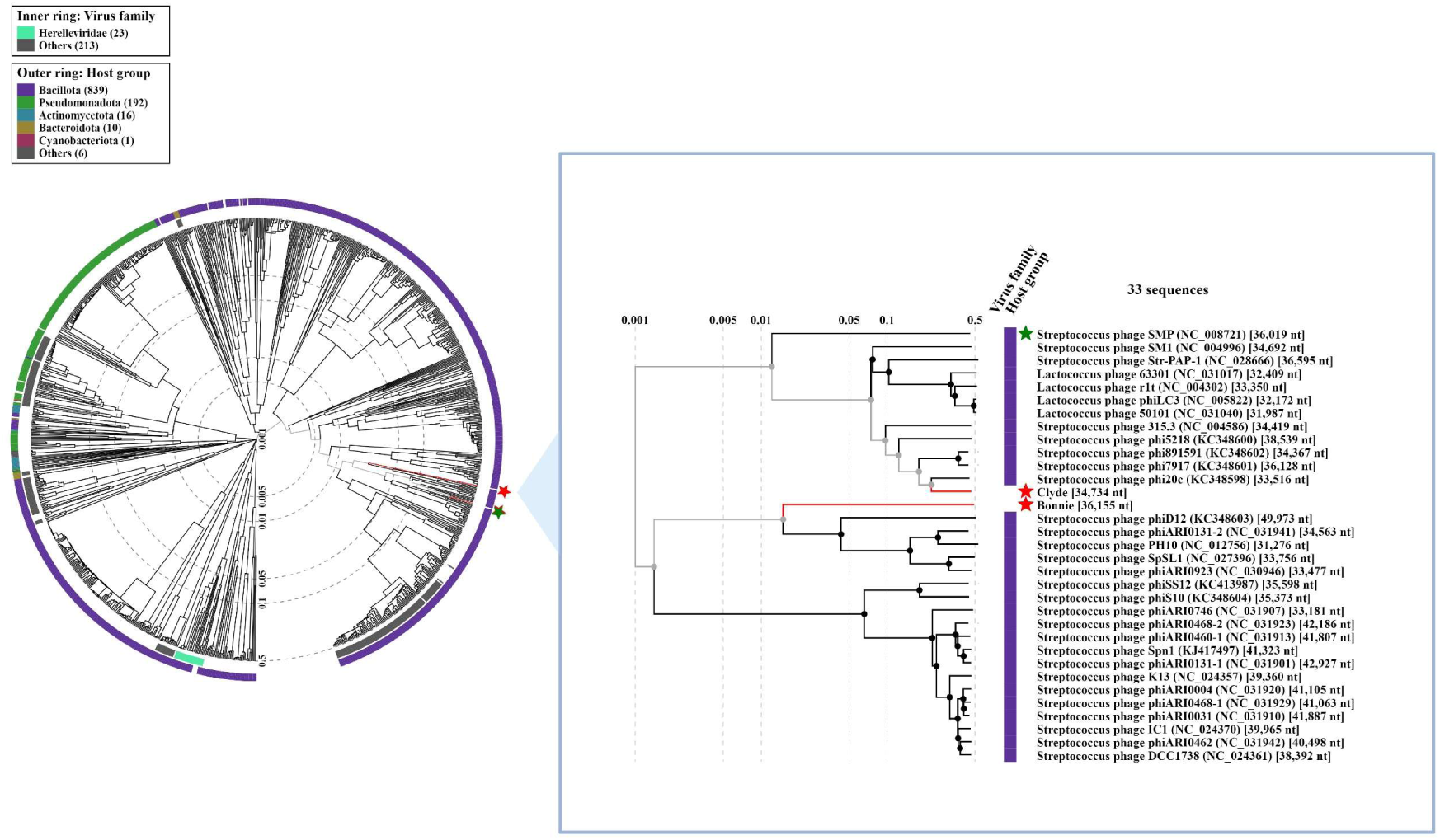
Viral proteomic tree of Bonnie and Clyde with related phages generated in ViPtree. Inner and outer rings represent ICTV virus family and host group. An expanded view of the tree shows Bonnie and Clyde (red) and the virulent phage SMP (green) with other closely related phages. The log scale bar (with dashed lines) represents the genomic similarity scores (SG) computed through normalised tBLASTx scores.

Furthermore, searches against the NCBI core nucleotide database using BLASTn did not identify any cultured phage with significant homology to Bonnie and Clyde. However, some deposited prophage sequences and regions in bacterial genomes (*S. suis*) shared some nucleotide similarity with Bonnie and Clyde (with at least ≥50% query coverage). Then together with the deposited prophage sequences and phage SMP, the intergenomic similarities were estimated in VIRIDIC. Three redundant prophages extracted from strains ISU2660, 90–1330, and DNR43 were subsequently removed as they were identical to a prophage in strain MA8 resulting in a total of 28 (pro)phages. The closest relative of Clyde is *Streptococcus* phage phi20c, a prophage from *S. suis* strain 8067, with a 52.4% intergenomic similarity. Bonnie shared 75.8% similarity with a predicted prophage from strain NLS50, which was isolated from a pig in the Netherlands in 2017 (Fig. 4A). The closest deposited prophage sequence to Bonnie is *Streptococcus* phage Javan597, which shares 61.8% intergenomic similarity. Furthermore, Clyde shares only 0.4% intergenomic similarity with Bonnie and 16.5% with phage SMP which is higher than Bonnie shares with SMP (4.4%).

**Fig. 4:**
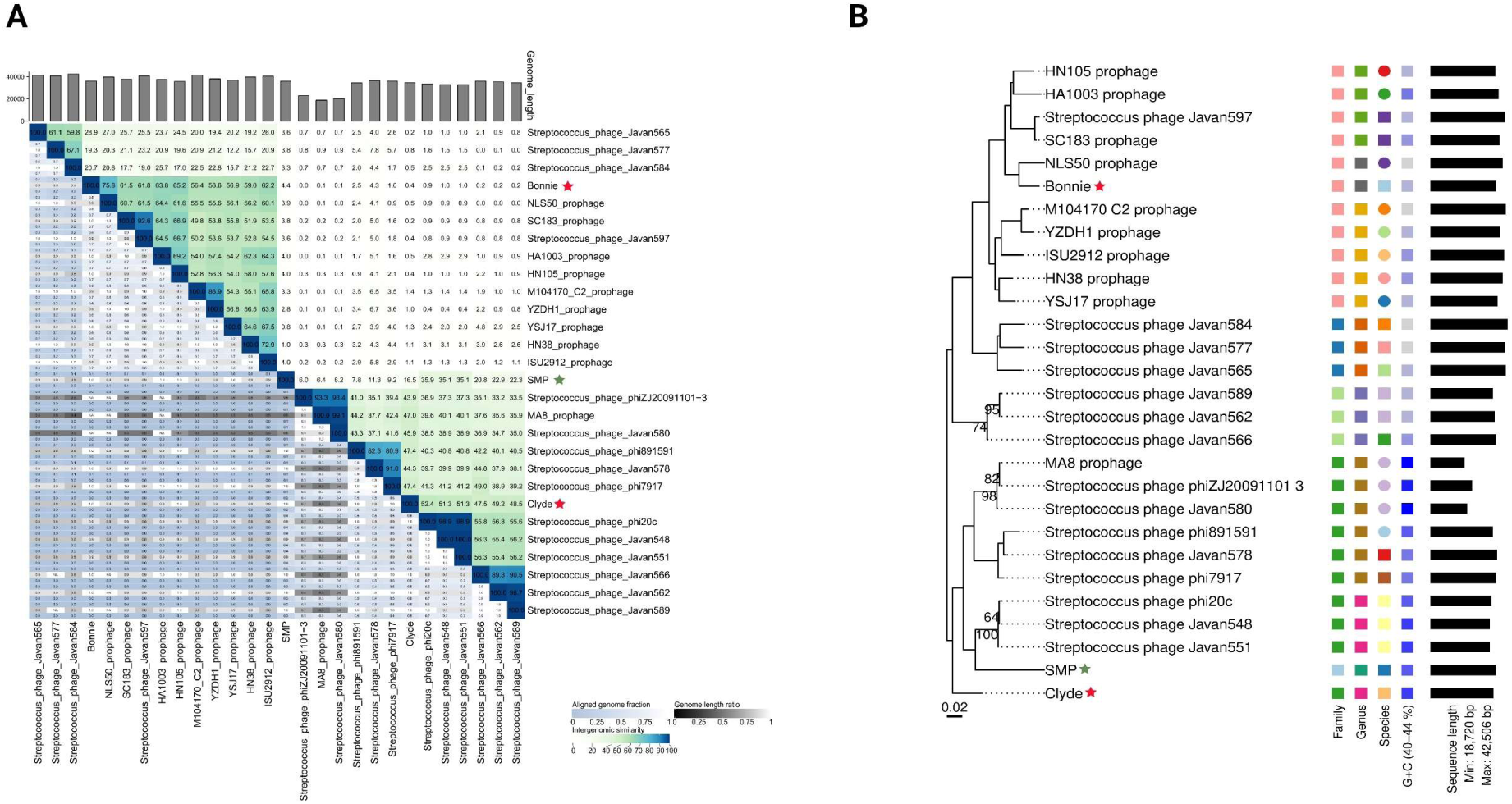
Phylogenomic analysis of Bonnie and Clyde. **(A)** Heatmap showing intergenomic similarities among Bonnie and Clyde, and closely related (pro)phages. (**B)** Phylogenomic GBDP tree of isolated phages inferred using formula D4 in VICTOR. Numbers above nodes (given that branch support exceeds ≥50%; nodes without annotated values indicate branches with lower support) represent pseudo-bootstrap support values from 100 replications. Tree was rooted at midpoint and the branch lengths are scaled using the GBDP distance formula *d_4_*. The scale bar (0.02) represents normalised dissimilarity between genomes. Coloured annotations on the right indicate the taxonomic clustering of phages based on ICTV cut-offs (Table S6). For each taxonomic rank, phages of the same taxon are assigned the same colour and/or shape. GC content is represented by gradient-coloured squares from ∼40% (min; grey) to ∼44% (max; blue) Genome length is represented by black horizontal bars.

Thus, based on ICTV genus (≥ 70% nucleotide identity) and species (≥ 95% nucleotide identity) delineation for dsDNA bacterial and archaeal viruses, phage Clyde represents a member of a novel species in a novel genus. Similarly, phage Bonnie belongs to a different novel species within a novel genus that includes the uncultivated prophage from strain NLS50. Additional phylogenomic analysis was conducted using VICTOR to generate a tree from the nucleotide sequence of the 28 (pro)phages to infer evolutionary relationships among the phages. This comparison yielded eight genus clusters spread across twenty-two species (Table S6). As predicted above, Bonnie and NLS50_prophage fall into two different species within a genus cluster (8). However, Clyde falls within a genus cluster (4) with its closest relative, phage phi20c, as well as *Streptococcus* phage Javan548 and Javan551. Phage SMP is the sole representative of its genus and species clusters (Fig. 4B).

The genome composition of Bonnie and Clyde and their closest relatives was analysed, including a predicted prophage (NLS50_prophage) and a deposited prophage (Javan597) for Bonnie, as well as phi20c for Clyde, with SMP used as a reference. Analysis with VirClust predicted 196 protein clusters among the six phages and each phage encoded 6 to 22 unique proteins (Table S7 and S8), however, no protein clusters were shared by all the six phages. One gene and three other genes were shared by five and four phages, respectively. Apart from Clyde, the other five phages encoded a conserved single-stranded DNA binding protein (SSB) with an average amino acid similarity of 84.8%. Other genes shared by four of the six phages included N-acetylmuramoyl-L-alanine amidase (84.3%), tail tape measure protein (58.8%), and an unknown protein (38.7%) (Table S9). Comparative genome alignment revealed high level of synteny particularly in the gene order of the head & packaging, as well as the tail modules. While this synteny was observed in all six phages, the average homology in these modules is low and conserved only among closest relatives (Fig. 5).

**Fig. 5:**
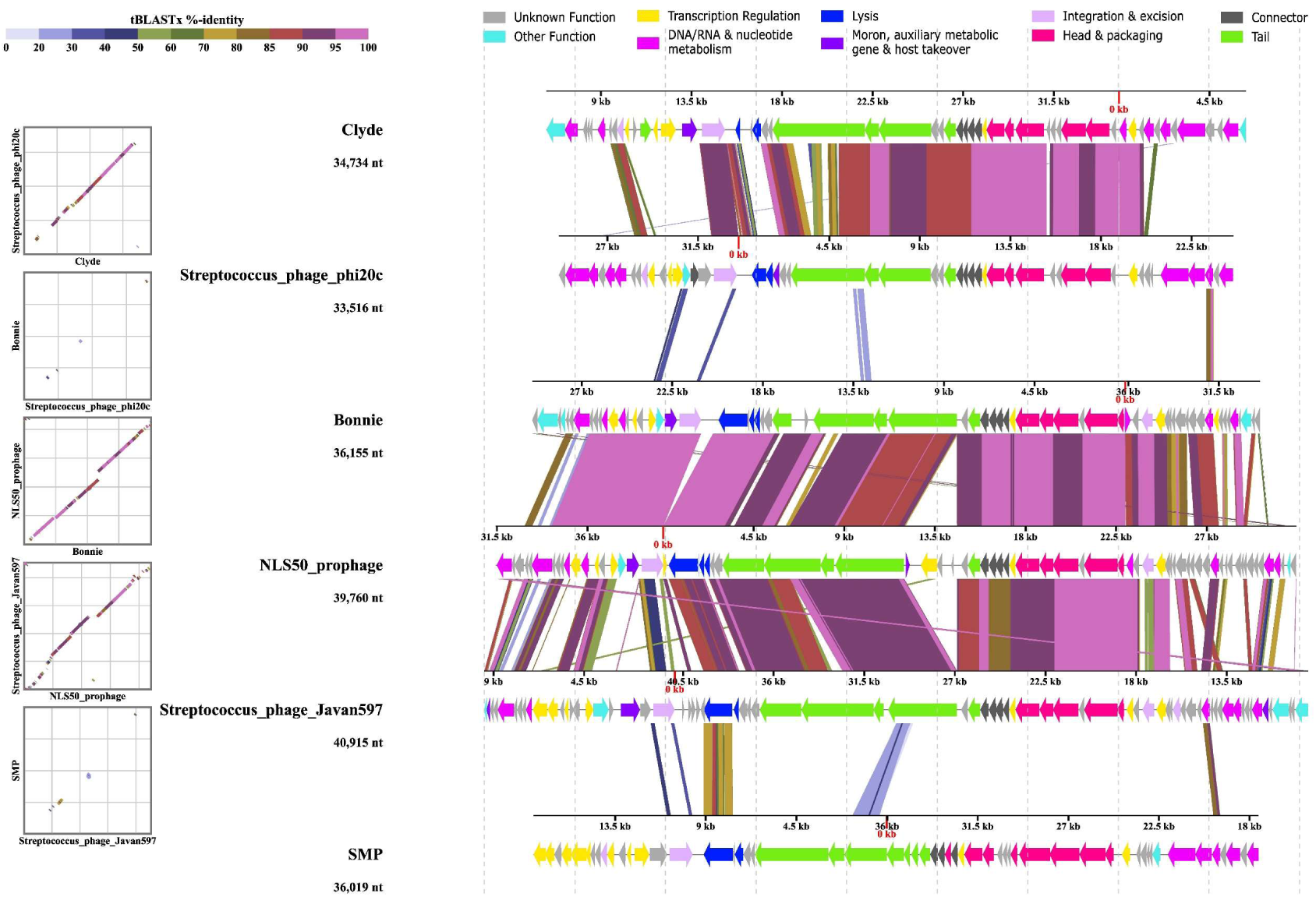
Comparative linear alignment displaying genomic features of isolated phages and their closest relatives. SMP is used as a reference. Arrows indicate position and direction of CDSs in genomes of the (pro)phages with colour-coding representing functional categories. Shaded regions between genomes represent levels of similarity computed with tBLASTx. Dot plot of pairwise genome alignments is shown on the far left, with a corresponding colour scale for both dot plot and alignments displayed in the top left. Both dot plot and alignments were generated in ViPTree using concatenated nucleotide sequence of the (pro)phages.

Apart from using whole phages in bacterial control, phage lysins have been characterised and purified for use against pathogens. The bactericidal activity of *S. suis* (pro)phage-derived lysins such as LySMP, Csl2, Ly7917, PlySs2 and PlySs9, as well as holins like HolSMP, has been experimentally validated both *in vitro* and *in vivo* against *S. suis* and other pathogens in previous studies (23, 24, 52–54). The protein sequence of LySMP (SMP) endolysin (481 amino acids) was used as reference for comparative analysis with lysins encoded by Bonnie, NLS50, Clyde, phi20c, and Javan597. These five phages encode lysins ranging from 247 to 468 amino acids in length and share between 19.3% to 79.3% amino acid similarity with the SMP endolysin (Fig. 6A and 6B). Pairwise amino acid identity matrix is supplied in Table S10.

**Fig. 6:**
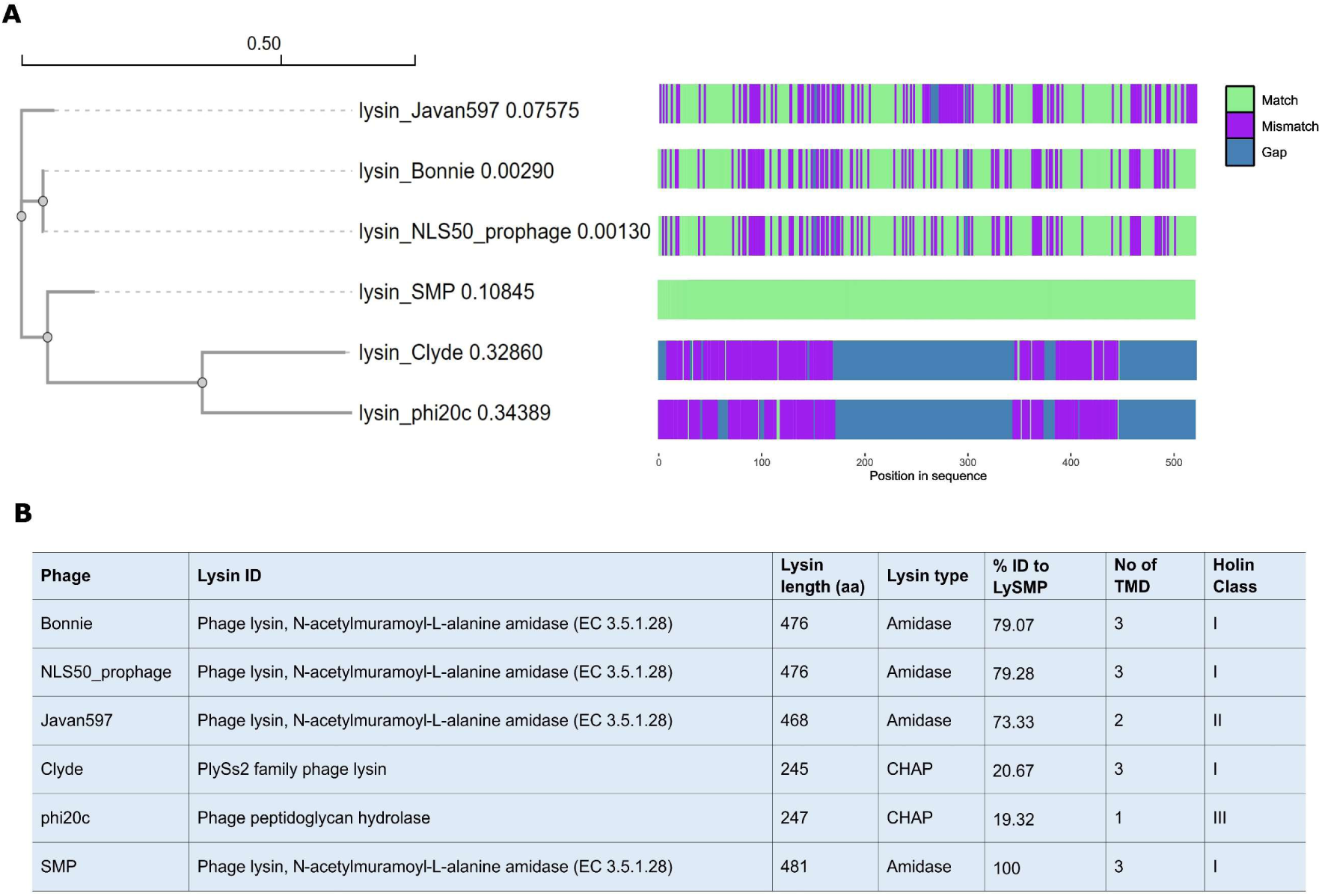
Phylogenetic tree of endolysins of the six (pro)phages. **(A)** Neighbour-joining tree based on amino acid sequence of endolysins constructed with Clustal Omega. Alignment was visualised with seqvisr tool using experimentally validated phage SMP endolysin (LySMP) as reference. (**B)** Predicted holin classes and endolysins. Holin transmembrane domains (TMDs) were predicted with DeepTMHMM. Lysin type is based on catalytic domains; N-terminal N-acetylmuramoyl-L-alanine amidase (amidase) and N-terminal cysteine/histidine-dependent amidohydrolase/peptidase (CHAP). Abbreviation: “aa” represents length in amino acids.

### Protein structure predictions and topological model assembly

The means by which *S. suis* phages recognises and bind to their host is currently unknown. To this end, we sought to predict the structure of the tail-associated proteins of Bonnie and Clyde including the predicted distal tail protein (Dit), tail-associated lysin (Tal), and, when present, the receptor-binding protein (RBP). We then compared the structure predictions of the adhesion machineries of Bonnie and Clyde, with those of previously isolated *S. suis* phage SMP, as well as the well characterised tail structure of *Streptococcus thermophilus* (St) phage STP1 (55) for which the host recognition and binding is described. In lambdoid tailed phages, the genes encoding the adhesion machinery are located between those encoding the tail tape measure protein (TMP) and the holin and lysin (56). The adhesion machineries typically comprise three main proteins: the Dit, the Tal, and, frequently, an RBP, which may occur in variable order (56). In St phages including STP1, these three genes are arranged sequentially in the Dit, Tal, and RBP order (55, 57). In Bonnie, all three genes were identified, whereas for Clyde and SMP, only the genes encoding Dit and Tal were present. Structural predictions were therefore made for the Dit and Tal gene products and their complexes, as well as for the RBP of Bonnie.

Dit proteins can be divided into two domains corresponding to the N- and C-terminal regions of the polypeptide chain. The N-terminal domain, known as the belt, comprises two β-sheets, a β-hairpin, and an α-helix (58, 59). The C-terminal domain called the galectin, is a two β-sheet structure similar to a galectin domain but lacking its saccharide-binding residues. The Dit protein of some phages, including that of phage Lambda, lack a galectin domain, while in others, such as phage T5, it is replaced by the OB-fold domain (60). Dits often contain carbohydrate-binding module (CBM) insertions within the galectin domain and are referred to as “evolved Dits” (61, 62). Evolved Dit proteins have been observed, as seen in the ∼500 amino acid-long Dit proteins of St phages, which also contain a CBM inserted within the galectin domain (55, 57). In contrast to STP1 (56) and other St phages, the Dits of Bonnie, Clyde, and SMP are not evolved and belong to the classical Dit family. As such, these Dits are unlikely to participate in host binding (Fig. 7A).

**Fig. 7:**
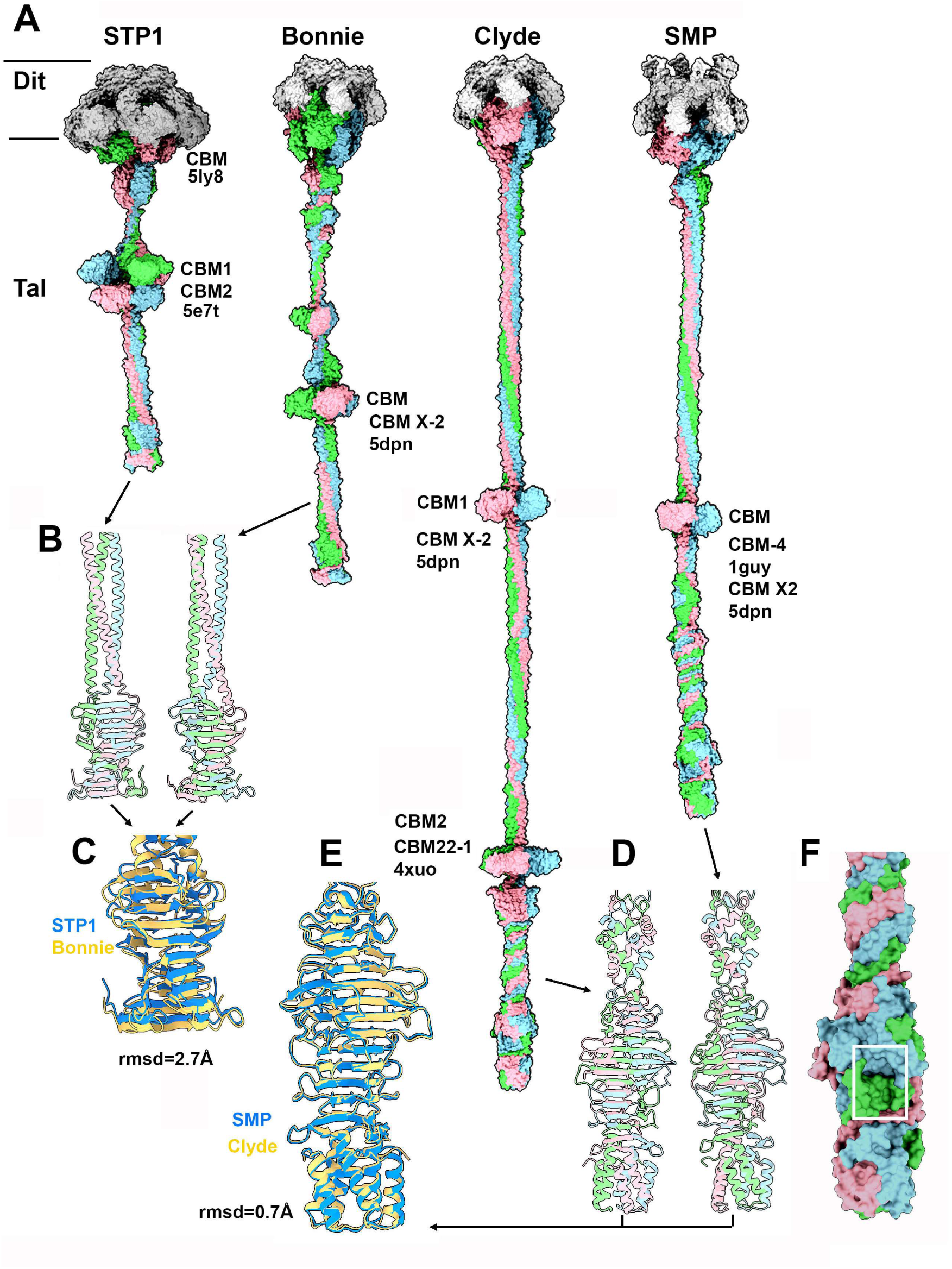
Structural representation of the Dits and Tals from *S. thermophilus* phage STP1 and *S. suis* phages Bonnie, Clyde and SMP. **(A)** Surface representation of the complexes formed by hexameric Dits and trimeric Tals. CBMs are annotated with their closest structural homologues. **(B)** Ribbon view of the Tal C-termini of STP1 and Bonnie. **(C)** Ribbon view of the superimposition of the Tal C-termini of STP1 (blue) and Bonnie (yellow). **(D)** Ribbon view of the Tal C-termini of Clyde and SMP. **(E)** Ribbon view of the superimposition of the Tal C-termini of SMP (blue) and Clyde (yellow). **(F)** Close-up surface view of the Tal C-terminus of Clyde. The crevice and putative receptor binding site is boxed (white).

The Tals of siphophages are trimeric and consist of an N-terminal structural domain of ∼350-400 amino acids (59, 63, 64). In many phages, this domain is followed by an extension that is believed to play a role in cell wall polysaccharide/peptidoglycan degradation, as seen in *L. lactis* P335 phage TP901-1 (64, 65), or in host binding as in the *Bacillus subtilis* phage SPP1 (66). The Tal extension of STP1 is 1,092 amino acids long, containing an N-terminal structural domain and two CBMs in the extension (Fig. 7A, STP1).

Bonnie possesses a trimeric Tal of 990 amino acids, with a classical N-terminal structural domain (residues 1-384). Unlike STP1, which possesses an Ig-like domain, Bonnie presents a tandem of small β-stranded domains (55)(Fig. 7A, Bonnie). This is followed by a triple β-stranded domain, forming a distinct small module, though no relevant similar structure was identified by Foldseek in the PDB. Short collagen linkers connect to a domain similar to the three β-domains described in the St phages’ Tals adjacent to a CBM. Foldseek identified this CBM as a member of the CBM X-2 family, according to the CAZy classification. After the CBM, a helical triplex abuts a β-prism as observed for STP1 (55).

In addition to Dit and Tal, Bonnie possesses RBPs. Structural predictions of the RBPs of Bonnie and STP1 as monomers identified a linear assembly of β-stranded domains forming a β-sandwich along with a complex C-terminal domain (Fig. 8A and B). Attempting to predict the full-length RBPs as a trimer revealed that the four β-sandwiches do not pack together, while the C-terminal domain forms a compact trimer (Fig. 8C and D; Fig. S1). The structure includes a small β-prism composed of 3×4 β-strands, followed by three packed β-sheets and terminated in three packed β-sandwiches. This last domain resembles the RBP heads of lactococcal phages, and Foldseek identified hits with RBP of the lactococcal phages TP901-1 and p2 (56, 67–69). While the RBP heads of STP1 and Bonnie exhibit similar folds, differences in the length of their loops and their sequences suggest they may target distinct polysaccharidic receptors (Fig. S2A).

**Fig. 8:**
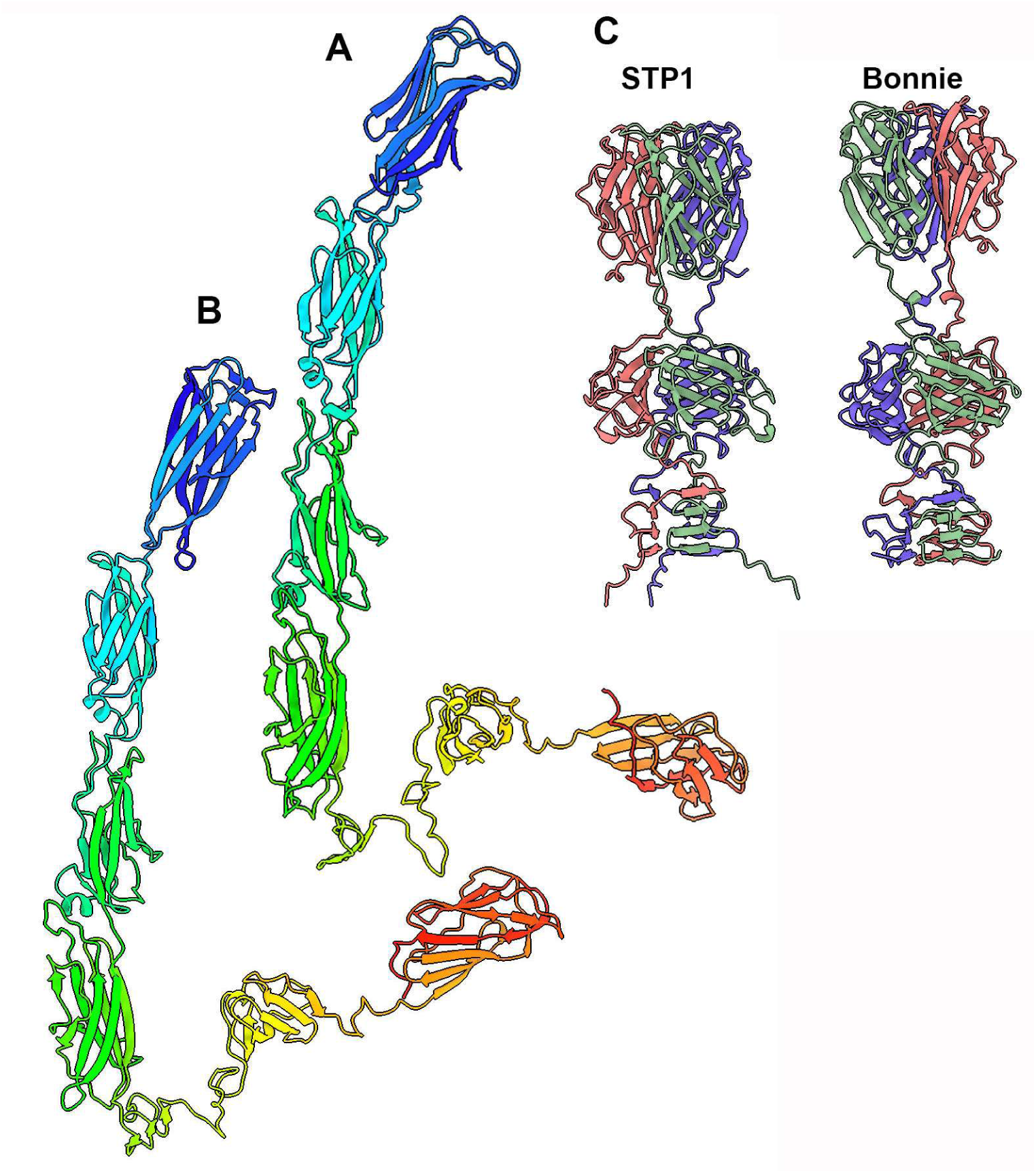
Structural representation of the RBPs from *S. thermophilus* phage STP1 and *S. suis* phage Bonnie. Ribbon representation of the RBP monomer from STP1 **(A)** and Bonnie **(B)**, rainbow coloured from its N-(blue) to C-terminus (red). **(C)** Ribbon representations of the RBP C-terminal trimer assemblies from STP1 and Bonnie.

Following the Dit hexamer, Clyde’s Tal N-terminal structural domain is immediately followed by a long triple helix and a CBM, which was identified by Foldseek as belonging to the CAZy CBM X-2 family (Fig. 7, Clyde). From this CBM, a long triple helix abuts another CBM, which is identified as belonging to the CAZy CBM22-1 family. Immediately following this, a large β-prism domain is observed, continuing into a complex triplex domain formed of β-turns and α-helices, which abuts a complex C-terminal domain (Fig. 7, Clyde). This C-terminal domain forms an intertwined β-prism, comprising 14 β-strands of variable length, decorated with long loops, and culminating in a C-terminal segment of two α-helices per monomer, forming a trimeric bundle of six α-helices (Fig. 7D and E). Foldseek retrieved a weak hit with the fibre tip of *Bdellovibrio bacteriovorus* H100 (Prob. 0.21; E-value 9.8×10^-1^; PDB ID: 8ond) (70). The *B. bacteriovorus* H100 fibre trimeric domain exhibits a large crevice at the interface between its monomers, a feature also observed in *Salmonella* phage P22, where it is filled by an O-antigen oligosaccharide (PDB ID: 30th0) (71). Notably, a similar crevice is also observed in Clyde’s Tal C-terminal domains (Fig. 7F), strongly suggesting that it may act as a binding domain for polysaccharidic receptors.

SMP exhibits very similar features when compared to Clyde, with the main difference being the shorter length of the second triplex helix and the presence of only one CBM, which is structurally similar to the CBM1 of Clyde (Fig. 7, SMP). The C-terminus of the Tal is highly superimposable to that of Clyde, as both Tal sequences share 98.4% identity in their last 250 residues (Fig. 7E, Fig. S2B)

### Plaque and virion morphology of Bonnie and Clyde

Both phages were propagated on strain 21171_DNR38 in THB. Bonnie produced clear, medium-sized plaques with frayed edges that measured 1.02 ± 0.05 mm in diameter and Clyde produced clear pinhead plaques measuring 0.29 ± 0.06 mm (Fig. 9A and 9C). TEM analysis revealed both phages exhibit a long, flexible, non-contractile tail and an icosahedral capsid. Clyde has a shorter tail compared to Bonnie, however, no obvious structures such as central fibre were observed at the base of either phage (Fig. 9B and 9D).

**Fig. 9:**
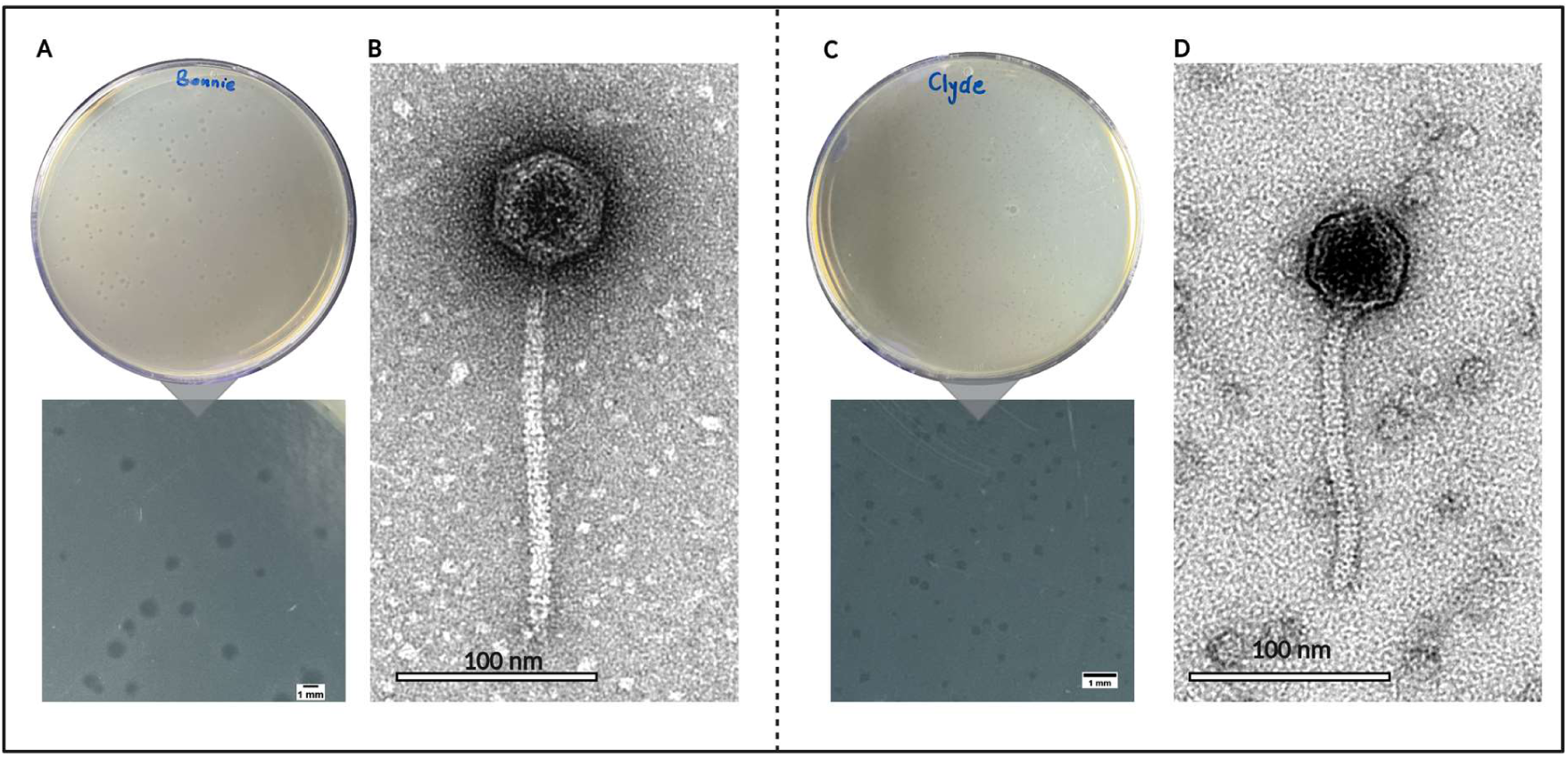
Plaque and virion morphology of Bonnie and Clyde. **(A)** plaques of Bonnie and **(C)** Clyde on double-layer agar plates. Plaque diameters of Bonnie (1.02 ± 0.05 mm) and Clyde (0.28 ± 0.06 mm) were measured with ImageJ software (National Institute of Health, Bethesda, USA). Scale bar represents 1 mm. Representative electron micrographs of **(B)** Bonnie and **(D)** Clyde. Bonnie has a capsid (56.27 ± 2.33 nm in diameter) and a long non-contractile tail of length 186.92 ± 5.92 nm and width 9.38 ± 0.81 nm. Clyde has a capsid that measures 54.22 ± 3.37 nm in length with a short non-contractile tail of 143.01 ±6.26nm and 10.2 ± 0.44 nm. Scale bar represents 100 nm.

### Host spectrum of Bonnie and Clyde

The susceptibility of 100 pathogenic *S. suis* strains to phages Bonnie and Clyde was evaluated by plaque assays. Bonnie produced plaques or clear zones on 15% of tested strains while Clyde formed plaques/clear zones on 58%. Collectively, the phages could infect 58 of the 100 strains tested. These susceptible strains include seven serotypes and thirteen known *S. suis* sequence types. Neither Bonnie nor Clyde could infect strains of other species (*S. thermophilus*, *L. cremoris*, and *E. coli*; Table S1). The EOP relative to strain 21171_DNR38 for 29 susceptible strains that formed plaques was tested. The EOP ranged from 0.001 to 1.2 (10^5^ to 10^8^ PFU/ml). Both Bonnie and Clyde produced plaques at a comparable efficiency on strains D71 and 21169_DNR36.

**Table 3:** Host range of Bonnie and Clyde.

| Strain | Bonnie | Clyde | Serotype | ST | Year | Pathotype | Country |
| --- | --- | --- | --- | --- | --- | --- | --- |
| 94_8576_4887 | - | + | 14 | 28 | 1994 | Respiratory | Canada |
| 89_5046 | - | + | 2 | 25 | 1989 | Systemic | Canada |
| 89_6891_2 | - | + | 2 | 25 | 1989 | Respiratory | Canada |
| 90_2741_7 | - | + | 2 | 25 | 1990 | Systemic | Canada |
| 1602951 | - | + | 2 | 25 | 2014 | Respiratory | Canada |
| 1596201 | - | + | 14 | 1 | 2014 | Systemic | Canada |
| 1463646 | - | + | 14 | 1 | 2013 | Systemic | Canada |
| 1667796 | - | + | 2 | NF | 2014 | Systemic | Canada |
| 1093404 | - | + | 27 | NF | 2008 | Systemic | Canada |
| 21137_DNR4 | - | + | 2 | 28 | 2018 | Respiratory | Denmark |
| 21136_DNR3 | - | + | 2 | 28 | 2018 | Respiratory | Denmark |
| 21135_DNR2 | - | + | 2 | 28 | 2018 | Respiratory | Denmark |
| 21169_DNR36 | + | + | 2 | 28 | 2018 | Respiratory | Denmark |
| 21171_DNR38 | + | + | 2 | 1 | 2018 | Respiratory | Denmark |
| 21172_DNR39 | + | + | 2 | 1 | 2018 | Respiratory | Denmark |
| 21173_DNR40 | - | + | 2 | 28 | 2018 | Respiratory | Denmark |
| 21176_DNR43 | - | + | 2 | 28 | 2018 | Respiratory | Denmark |
| D1 | - | + | 2 | 1 | 2019 | Unknown | Ireland |
| D3 | + | + | 2 | 1 | 2019 | Respiratory | Ireland |
| D5 | - | + | 2 | 1 | 2019 | Systemic | Ireland |
| D6 | + | + | 2 | 1 | 2019 | Systemic | Ireland |
| D7 | + | + | 2 | 1 | 2019 | Unknown | Ireland |
| D8 | - | + | 2 | 28 | 2012 | Respiratory | Ireland |
| D10 | - | + | 2 | 1 | 2012 | Systemic | Ireland |
| D14 | - | + | 2 | 1 | 2011 | Systemic | Ireland |
| D16 | - | + | 14 | 1 | 2011 | Unknown | Ireland |
| D17 | + | + | 2 | 1 | 2011 | Systemic | Ireland |
| D19 | - | + | 2 | 1 | 2010 | Respiratory | Ireland |
| D23 | - | + | 14 | 1 | 2008 | Unknown | Ireland |
| D24 | - | + | 2 | 1 | 2012 | Respiratory | Ireland |
| D29 | - | + | 7 | 29 | 2008 | Respiratory | Ireland |
| D31 | - | + | 4 | 856 | 2008 | Unknown | Ireland |
| D32 | + | + | 2 | 2629 | 2007 | Unknown | Ireland |
| D35 | - | + | 2 | 2646 | 2019 | Respiratory | Ireland |
| D42 | - | + | 16 | 2632 | 2019 | Respiratory | Ireland |
| D46 | + | + | 2 | 1 | 2019 | Respiratory | Ireland |
| D49 | - | + | 16 | 2632 | 2019 | Systemic | Ireland |
| D52 | + | + | 2 | 1 | 2018 | Systemic | Ireland |
| D57 | - | + | 2 | 2646 | 2019 | Respiratory | Ireland |
| D58 | + | + | 14 | 1 | 2019 | Systemic | Ireland |
| D68 | + | + | 2 | 1 | 2020 | Unknown | Ireland |
| D71 | + | + | 14 | 1 | 2020 | Systemic | Ireland |
| D75 | - | + | 9 | 2640 | 2020 | Respiratory | Ireland |
| D78 | - | + | 2 | 1 | 2020 | Respiratory | Ireland |
| D79 | - | + | 2 | 25 | 2020 | Systemic | Ireland |
| D84 | - | + | 2 | 2641 | 2021 | Unknown | Ireland |
| D93 | - | + | 2 | 28 | 2021 | Systemic | Ireland |
| D94 | - | + | 14 | 124 | 2021 | Systemic | Ireland |
| 19858_M105281_R30 | - | + | 7 | 24 | 2017 | Respiratory | Spain |
| 19867_M106485_R39 | + | + | 2 | 1 | 2018 | Respiratory | Spain |
| 19779_M101513_S1 | - | + | 2 | 1 | 2017 | Systemic | Spain |
| 19797_M104300_S19 | - | + | 2 | 1 | 2017 | Systemic | Spain |
| 19798_M104300_S20 | - | + | 2 | 1 | 2017 | Systemic | Spain |
| 19802_M105040_S24 | - | + | 2 | 1 | 2017 | Systemic | Spain |
| 19803_M105040_S25 | - | + | 2 | 1 | 2017 | Systemic | Spain |
| 19806_M105244_S28 | - | + | 2 | 1 | 2017 | Systemic | Spain |
| 19807_M105244_S29 | - | + | 2 | 1 | 2017 | Systemic | Spain |
| 19785_M102095_S7 | + | + | 2 | 1 | 2017 | Systemic | Spain |

**Fig. 10:**
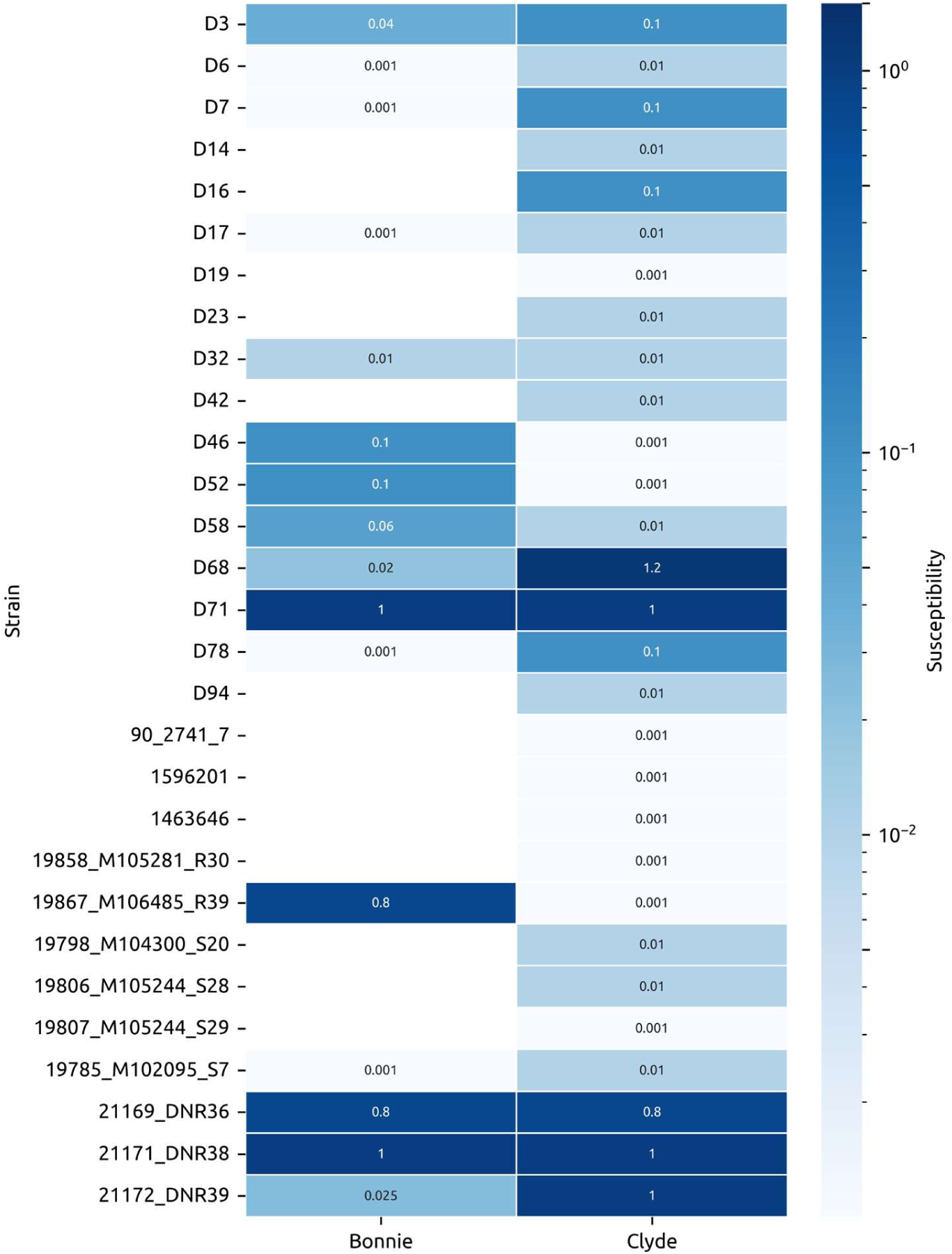
Efficiency of plating. The EOP relative to strain 21171_DNR38 was estimated using spot assays for 29 susceptible strains that formed plaques in host range analysis. The EOP values ranged from 0.001 to 1.2 (10^5^ to 10^8^ PFU/ml) for strains with detectable plaque formation. Strains where no plaques were observed (EOP = 0.00) are indicated by blank white boxes.

### *In vitro* lytic activity of Bonnie and Clyde

The changes in bacterial density in response to phage exposure was monitored over 24 hours. On strain 21171_DNR38, the bactericidal activity of Clyde was generally proportional to the MOI in a dose-dependent manner. All MOIs tested showed a statistically significant difference compared to the no-phage group (*p* < 0.0001). However, at higher MOIs (100 and 10), regrowth of resistant populations was observed at 8 hours and 20 hours post-infection, respectively (Fig. 11B). In the case of Bonnie, highest inhibition was observed at MOI 10 (highest MOI tested) and MOI 0.001 (*p* < 0.0001). Near the 6-hour time point, an exponential increase in bacterial density, comparable to the no phage control was recorded at MOI 1, 0.1 and 0.01 (Fig. 11A). Upon infection with a cocktail of Bonnie and Clyde, 21171_DNR38 growth was inhibited, particularly at MOIs 10 and 1 for up to 6 hours, following which bacterial density increased exponentially. Based on Clyde’s lytic activity at MOI 10, two different mixed-strain cultures were exposed to Clyde at this MOI. The growth of both multi-strain mix A (21171_DNR38, 19867_M106485_R39, DNR36, D71, and M105040_S24) and multi-strain mix B (21171_DNR38, D94, D8, D52, and D75) was significantly supressed (*p* < 0.0001), although not completely.

**Fig. 11:**
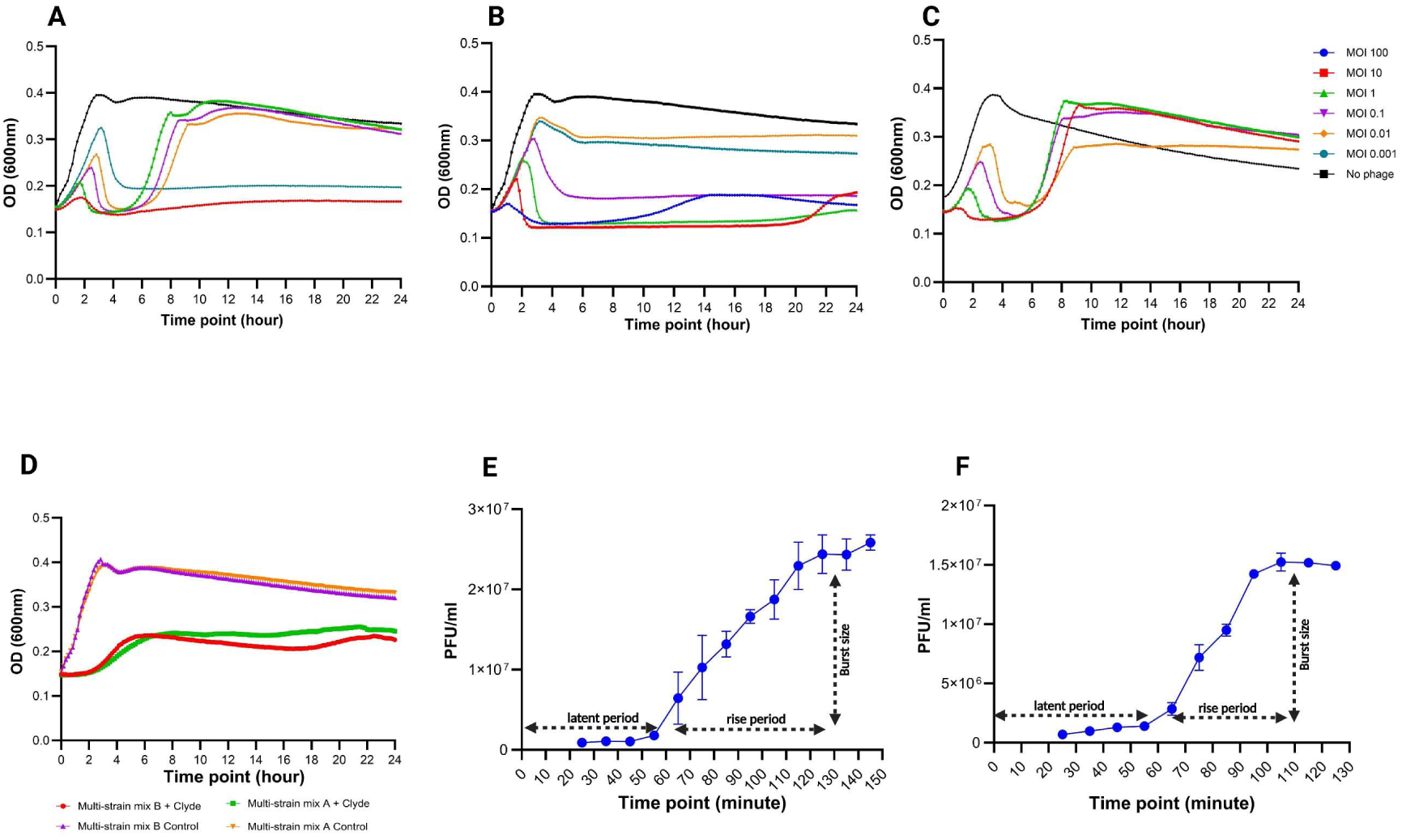
*In vitro* lytic activity and one-step growth curves. Bacterial killing activity of **(A)** Bonnie, **(B)** Clyde and **(C)** a cocktail of both phages at different MOIs was tested *in vitro*. OD_600_ was read every 10 minutes for 24 hours. **(D)** Lytic activity of Clyde (MOI 10) against two multi-strain cultures was monitored for 24 hours. Multi-strain mix A (21171_DNR38, 19867_M106485_R39, DNR36, D71, and M105040_S24) and Multi-strain mix B (21171_DNR38, D94, D8, D52, and D75). One step growth curve for **(E)** Bonnie and **(F)** Clyde at MOI 0.01. The error bars represent the standard error of the mean from independent replicate experiments.

### One step growth curve and burst size

One-step growth experiments determined Clyde had an approximate latent period of 55 minutes and a burst size of 35 PFU per infected cell. The latent period of Bonnie was estimated to be 55 minutes with a smaller burst size of 27 PFU per cell (Fig. 11E and 11F).

### Stability of Bonnie and Clyde under different pH and temperature conditions

The stability of phages under different thermal and pH conditions is crucial for storage and downstream processing. To evaluate this, the stability of Bonnie and Clyde was assessed by incubating aliquots at various temperatures and pH levels. For Clyde, titres remained stable at 37, 40 and 50°C for up to 120 minutes, showing no significant difference compared to storage temperature (4°C) (*p* > 0.05). However, at 60°C phage titres dropped below detection limits within 30 minutes. Titres of Bonnie did not significantly decrease at 37 or 40°C following exposure for up to 120 minutes compared to 4°C (*p* > 0.05). However, at 50°C, titres gradually declined, with a 3.2 log PFU/mL loss from the initial 8.6 log PFU/mL after 120 minutes. At 60°C, titres significantly dropped to 2.9 log PFU/mL by 30 minutes and by 90 minutes no plaques were detected. Phage stability was lost at 70°C for both phages. When incubated in pH-adjusted SM buffer (2,3,4,5,6,7,8,9,10,11,12 and 13) for 1, 2 or 24 hours, no significant change in viability was recorded across time points. There was complete inactivation of phage particles at pHs 2 and 13. Howover, both phages were stable from pH 4 to pH 10.

**Fig. 12:**
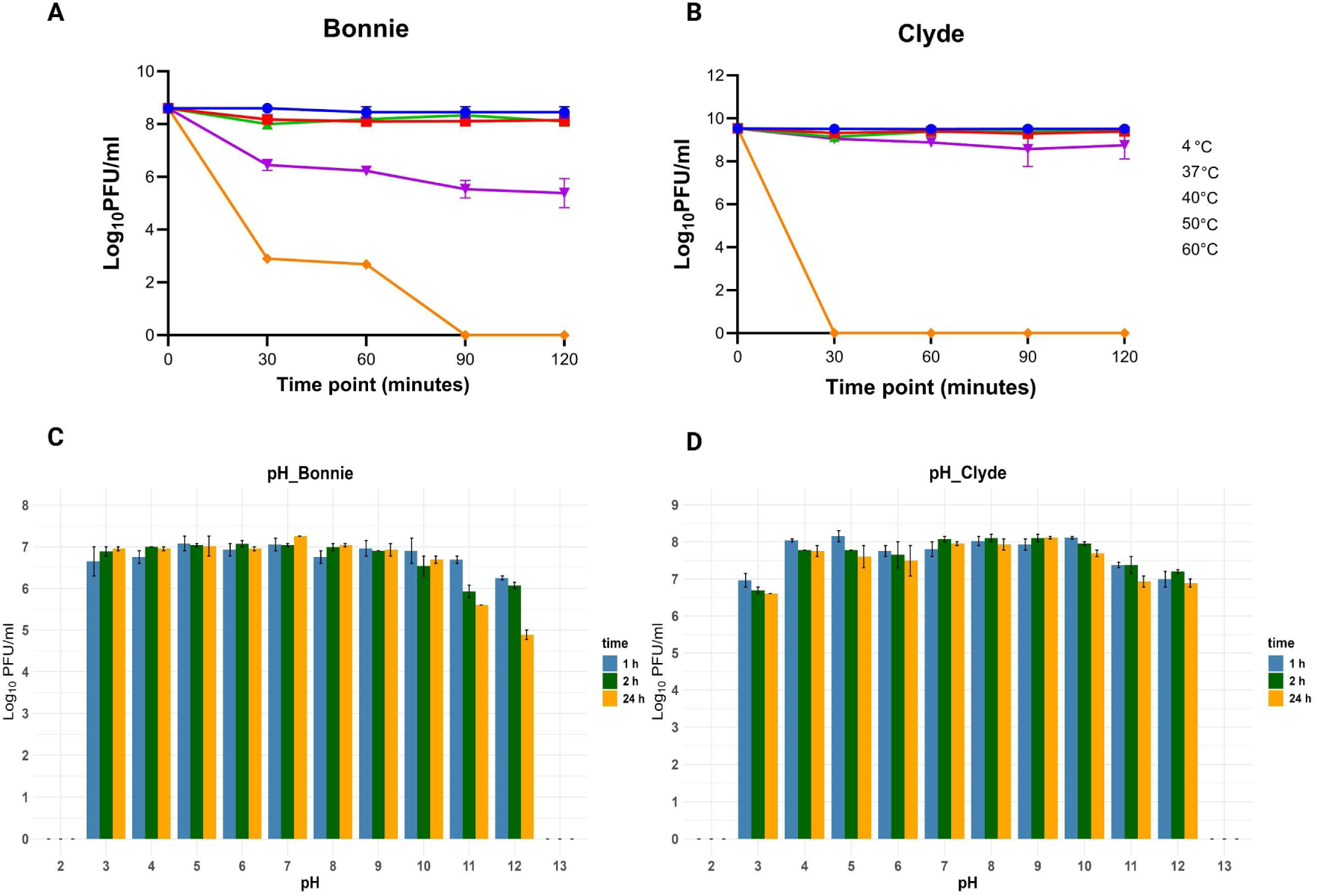
Stability of phages under different physicochemical conditions. Stability of Bonnie **(A)** and Clyde **(B)** at different temperatures was monitored over 120 minutes. Stability of Bonnie **(C)** and **(D)** Clyde was assessed following incubation in different pH-adjusted SM buffer for 1 hour, 2 hours, and 24 hours. Error bars represent the standard error of the mean from independent replicate experiments.

## Discussion

In this study, we report the isolation of two temperate phages that infect *S. suis*: Bonnie, isolated from pig tissue, and Clyde, obtained through prophage induction. Despite extensive screening of diverse samples, only Bonnie was isolated during the screening of pig-derived samples by plaque assay. Although the previously isolated *S. suis* phage SMP has been described as “lytic” and “virulent”, in our study, PhaTYP analysis predicted its lifestyle as temperate (Table S5). There are currently several bacteria for which virulent phages have not been reported, including *Legionella pneumophila*, *Clostridioides difficile*, *Bifidobacterium* species, and *S. suis* (72–74). A potential reason for the difficulty in isolating phages against *S. suis* is due to the high level of phase variation in the defence mechanisms of the bacterium (25, 75, 76). This would mean that in a culture derived from a single colony, phase variation rapidly produces genotypically and phenotypically diverse subpopulations, some of which cannot be infected by phage leading to their growth in a plaque assay, and a failure to observe plaques. We made a similar observation when Bonnie was passaged on 19867_M106485_R39—the initial isolation host—where plaque formation was inconsistent based on the version of bacterial stock used. Modulation of bacterial susceptibility to phage through phase variation has been described in other species including *Campylobacter jejuni*, *C. difficile*, and *Haemophilus influenzae* (77–79). In the case of *H. influenzae*, phage extinction was observed when the resistant subpopulation exceeded 34% (79). Future work will use 19867_M106485_R39 to investigate the mediation of phage susceptibility by phase-variable genes.

While MitC is routinely used to induce prophages in bacteria, previous attempts to induce prophages in *S. suis* lysogens harbouring full-length prophages were largely unsuccessful, with only two prophages induced from fifty-six isolates (80). All three methods used in this study successfully induced the expected prophage (Clyde) from 19867_M106485_R39; however, the resulting plaques were turbid and of low titres. Spiking the filtered induction lysates with 21171_DNR38 improved plaque formation and enabled subsequent single-plaque purification. As previously demonstrated in other bacteria (81–84), temperature cycling was shown to trigger prophage induction in *S. suis*

Genomic analysis revealed that Bonnie and Clyde shared only 4.4% and 16.5% nucleotide similarity with phage SMP, respectively. A search could not identify any closely related cultivated phage. Consequently, Bonnie and Clyde were compared to SMP, 15 publicly available *S. suis* prophages, and ten prophages predicted from publicly available host genomes, based on nucleotide similarity estimated using BLASTn. This comparison confirmed that Bonnie and Clyde are two novel species that belong to two distinct novel genera. Orthologous proteins from six (pro)phages, including Bonnie and Clyde, their closely related phages, and SMP as reference, were grouped into protein clusters. While no core clusters were identified, all (pro)phages, except Clyde, encode a conserved SSB protein, a protein previously identified in seven of twelve *S. suis* (pro)phages (80). The SSB bind to single-stranded DNA with high affinity to protect against degradation and formation of secondary structure during phage DNA replication (85). In its absence, phages co-opt host SSB for viral replication. The use of SSB by phages is so common that they have recently been shown to activate defence systems such as Hna, Retron-Eco8, Hachiman, and AbpAB (86, 87). The previously characterised PlySs2, which has a CHAP domain shares 90.2% protein identity with the endolysin of Clyde (23). In contrast, the endolysin encoded by Bonnie, and the other *S. suis* phages is a N-acetylmuramoyl-L-alanine amidase. Lysins and holins of *S. suis* phages have previously been purified and tested both *in vitro* and *in vivo*. The endolysins of Bonnie and Clyde could further be developed and evaluated for their specific antimicrobial activity against *S. suis*.

Structural predictions of the adhesion device proteins of Bonnie and Clyde revealed two distinct lineages of *S. suis* phages: St-like, exemplified by STP1 (55), and *S. suis*-like, similar to SMP. Bonnie’s structure aligns with the St phage-like lineage, possessing an RBP, while Clyde and SMP are specific to phages infecting *S. suis.* The *S. suis* lineage phages lack RBPs but display a Tal C-terminal domain that may function in receptor-binding. Regarding the Tal C-terminus of Clyde (or SMP), it is noteworthy that Foldseek returned many hits from AlphaFold database (AFDB) consisting of Tal C-termini from *S. suis* and other streptococcal phages, but not from St phages (Fig. S3A). Similarly, despite the overall similarity of Bonnie’s RBP C-terminus to those St phages, submitting Bonnie’s RBP C-terminus to Foldseek also returned hits in the AFDB from Tals of *S. suis* and other streptococcal phages, but not from St phages (Fig. S3B). These observations suggest that, from the perspective of adhesion device, two very different families of phages can specifically infect *S. suis*. Furthermore, despite the lack of nucleotide similarity and their different ecological niches, the structural similarities between the adhesion devices of Bonnie and St phages, might point to two possible evolutionary scenarios. The structural similarities could arise from a common ancestral gene, connecting *S. suis* and *S. thermophilus* through divergent evolution, as observed in the structurally similar RBP (68) and capsid proteins (88) shared by some mammalian viruses and phages. Alternatively, the observed similarities may be the result of convergent evolution, where these similar structures have evolved independently to adapt for adhesion to host surfaces. Such convergence has been described between the baseplates of coliphage T4 and lactococcal phage p2 (63), as well as the Dits of phages infecting *S. aureus*, *Bacillus subtilis,* and *L. lactococcus* (89). Both scenarios highlight potential complex evolutionary links between pathogenic and non-pathogenic streptococcal species.

Bonnie and Clyde have long, non-contractile tails consistent with previously reported *S. suis* (pro)phages Ss2, and SMP (19, 90). Collectively, Bonnie and Clyde infect 58/100 strains tested, with Bonnie displaying a narrower host range (15/100 strains), similar to SMP. Every strain susceptible to Bonnie was also susceptible to Clyde albeit with varying EOP. This observation is likely driven by transient factors—such as mutations in host receptors and acquisition of anti-phage systems through horizontal gene transfer—shaped by coevolutionary dynamics and evolutionary trade-offs within the same host species rather than fixed intrinsic barriers like non-host resistance due to differences in receptor binding sites observed in host range at higher taxonomic levels (91). Bonnie and Clyde’s differential host range may result from receptor-level adaptations, with Clyde’s adhesion device targeting a subset of receptors not recognised by Bonnie’s RBP. These specificities align with nested bipartite interaction networks, where specialist phages like Bonnie target subsets of generalist phages’ hosts (92, 93). Moreover, anti-phage defence systems may contribute to the observed host range. As we showed previously, *S. suis* genomes encode between one to ten of 20 distinct defence systems that may target different stages of the phage life cycle (20). Bonnie-resistant strains may harbour defence systems that are activated by Bonnie-specific triggers such as SSB, while Clyde overcomes these barriers through alternative infection strategies. Future transcriptomic and knockout studies could investigate these dynamics by exploring host responses to Bonnie and Clyde using Clyde-susceptible, Bonnie-resistant strains, as well as bacteriophage-insensitive mutants (BIMs). Bonnie and Clyde were first co-isolated when superinfection of 19867_M106485_R39 by Bonnie triggered the induction of Clyde (prophage). Such induction events increase the likelihood of recombination between the two phages which may lead to the emergence of new phage genotypes with wider host specificities (94, 95). Additionally, induced phages could infect and lyse cells resistant to the superinfecting phage (96) thereby potentially improving therapeutic outcomes.

We investigated the *in vitro* lytic activity of the phages against 21171_DNR38 at different MOIs. Clyde generally inhibited 21171_DNR38 growth in an MOI-dependent manner whereas with Bonnie, the highest inhibition was observed at MOI 10 and 0.001. Resistance to Bonnie and Clyde was observed at certain MOIs (Fig. 11A and 11B) by 6 hours and 8 hours, respectively. In the context of phage therapy, BIMs have been shown to display attenuated virulence *in vivo* (97), facilitate the reversion of antibiotic resistance (98), and sensitise resistant strains to other infecting phages (99). Phenotypic characterisation and *in vivo* experiments could be used to evaluate the fitness trade-offs in Bonnie and Clyde-resistant mutants.

Combining the phages into cocktails did not improve lytic activity beyond that observed with individual phages, as the activity of the cocktail was comparable to Bonnie alone, with resistance emerging by 6 hours post-infection in three of the five MOIs tested. While phage cocktails have been shown to improve bacterial clearance and minimise emergence of phage-resistant populations through synergistic interactions, antagonism is not uncommon, where similar phages compete for the same surface molecules (16, 100). Given that Bonnie and Clyde have different adhesion devices, antagonism seem less likely, but our results are more consistent with the findings in *E. coli* infecting-phages, where the efficacy of individual phages did not predict efficacy of cocktails (101). When tested *in vitro* against two multi-strain cultures that mimic the polyclonal nature of infection, Clyde significantly inhibited bacterial growth. Each multi-strain group (Fig. 11D) consisted of strains from two or three different serotypes (serotypes 2, 9, and 14), which are among the most commonly implicated agents in human and animal *S. suis* infections globally.

Phage stability is a key factor in their application as therapeutics, particularly as the acidic environment of the mammalian gastrointestinal tract can inactivate free phages when administered orally. Both Bonnie and Clyde were shown to be very stable across different pH and temperature conditions. Downstream processing such as encapsulation, freeze drying, and formulation into inhalable dry powders and steam pellet feed can be used to improve stability in different temperatures and extend shelf-life in phages such as Bonnie and Clyde (102, 103).

Overall, the phages described herein will support future investigations into the evolutionary relationships between phages and *S. suis* in the context of host recognition and infection, anti-phage defence mechanisms, and bacterial virulence. Beyond fundamental research, the demonstrated stability and *in vitro* activity of Bonnie and Clyde highlight their potential as antibacterial agents either through engineering obligately virulent mutants, or using their encoded proteins, such as endolysins.

## Supporting information

Table S4 to S10: In silico analysis

## Data availability

Coordinates of predicted structures are accessible on Zenodo (https://zenodo.org/records/14212611). Nucleotide sequences of phages Bonnie and Clyde were deposited in GenBank under accession numbers PQ720431 and PQ0432, respectively. Accession numbers of host genomes are presented in supplementary data.

## Supplementary materials

Figure S1 to S3 and Table S1 to S3: https://zenodo.org/records/14610606 Table S4 to S10: https://zenodo.org/records/14610606

## CRediT authorship contribution statement

**Emmanuel Kuffour Osei:** conceptualisation, data curation, formal analysis, investigation, methodology, visualisation, writing – original draft, writing – review and editing. **Reuben O’Hea:** data curation and investigation. **Christian Cambillau:** investigation, methodology, visualisation, software, writing – original draft, writing – review and editing. **Ankita Athalye:** data curation and investigation. **Frank Hille:** methodology, visualisation, software. **Charles M.A.P. Franz:** visualisation, software. **Áine O’Doherty:** data curation and investigation. **Margaret Wilson:** data curation and investigation. **Gemma G R Murray:** Writing – review and editing. **Lucy A Weinert:** data curation and investigation, writing – review and editing. **Edgar Garcia Manzanilla:** funding acquisition, methodology, writing – review and editing. **Jennifer Mahony and John G Kenny:** conceptualisation, funding acquisition and project management, investigation, methodology, writing – original draft, writing – review and editing.

## Acknowledgement

We would like to thank John Moriarty and the Pathology Division (Department of Agriculture Laboratories, Backweston) for generously providing some of the bacterial strains used in this study, as well as Marcelo Gottschalk and Nahuel Fittipaldi for the Canadian isolates. Transmission electron microscopy was carried out with technical assistance from Morten Bratschke (Max Rubner-Institut). We thank members of the Mahony Lab for assistance during the CsCl purification. We are grateful to Mário Ornelas for his assistance in sampling. We are grateful for the HPC resources from GENCI-IDRIS [2023-AD010714075; in part] and INRAE MIGALE bioinformatics facility. We acknowledge UCSF ChimeraX which is developed by the Resource for Biocomputing, Visualisation and Informatics at the University of California, San Francisco, with support from National Institutes of Health [R01-GM129325]. We are grateful to Gabriele A. Lugli and Marco Ventura (GenProbio s.r.l, Parma, Italy) for carrying out the genome sequencing of the phages. Authors are grateful to Ramya Balasubramanian and other members of Vision 1 Lab at Teagasc Moorepark for their insights and constructive feedback on the analysis and preparation of the manuscript.

## References

1. The emergence and diversification of a zoonotic pathogen from within the microbiota of intensively farmed pigs - PMC. https://pmc.ncbi.nlm.nih.gov/articles/PMC10666105/. Retrieved 9 November 2024.

2. Frontiers | Porcine respiratory disease complex: Dynamics of polymicrobial infections and management strategies after the introduction of the African swine fever. https://www.frontiersin.org/journals/veterinary-science/articles/10.3389/fvets.2022.1048861/full. Retrieved 9 November 2024.

3. Perch B, Kristjansen P, Skadhauge Kn. 1968. Group R Streptococci Pathogenic for Man. Acta Pathologica Microbiologica Scandinavica 74:69–76.

4. Rayanakorn A, Goh B-H, Lee L-H, Khan TM, Saokaew S. 2018. Risk factors for Streptococcus suis infection: A systematic review and meta-analysis. Sci Rep 8:13358.

5. Segura M, Fittipaldi N, Calzas C, Gottschalk M. 2017. Critical Streptococcus suis Virulence Factors: Are They All Really Critical? Trends in Microbiology 25:585–599.

6. Tien LHT, Nishibori T, Nishitani Y, Nomoto R, Osawa R. 2013. Reappraisal of the taxonomy of *Streptococcus suis* serotypes 20, 22, 26, and 33 based on DNA–DNA homology and *sodA* and *recN* phylogenies. Veterinary Microbiology 162:842–849.

7. Update on Streptococcus suis Research and Prevention in the Era of Antimicrobial Restriction: 4th International Workshop on S. suis - PMC. https://pmc.ncbi.nlm.nih.gov/articles/PMC7281350/. Retrieved 9 November 2024.

8. Weinert LA, Chaudhuri RR, Wang J, Peters SE, Corander J, Jombart T, Baig A, Howell KJ, Vehkala M, Välimäki N, Harris D, Chieu TTB, Van Vinh Chau N, Campbell J, Schultsz C, Parkhill J, Bentley SD, Langford PR, Rycroft AN, Wren BW, Farrar J, Baker S, Hoa NT, Holden MTG, Tucker AW, Maskell DJ. 2015. Genomic signatures of human and animal disease in the zoonotic pathogen Streptococcus suis. Nat Commun 6:6740.

9. Okura M, Auger J-P, Shibahara T, Goyette-Desjardins G, Van Calsteren M-R, Maruyama F, Kawai M, Osaki M, Segura M, Gottschalk M, Takamatsu D. 2021. Capsular polysaccharide switching in Streptococcus suis modulates host cell interactions and virulence. Sci Rep 11:6513.

10. Hadjirin NF, Miller EL, Murray GGR, Yen PLK, Phuc HD, Wileman TM, Hernandez-Garcia J, Williamson SM, Parkhill J, Maskell DJ, Zhou R, Fittipaldi N, Gottschalk M, Tucker AW(. D, Hoa NT, Welch JJ, Weinert LA. 2021. Large-scale genomic analysis of antimicrobial resistance in the zoonotic pathogen Streptococcus suis. BMC Biol 19:191.

11. Dechêne-Tempier M, de Boisséson C, Lucas P, Bougeard S, Libante V, Marois-Créhan C, Payot S. 2024. Virulence genes, resistome and mobilome of Streptococcus suis strains isolated in France. Microbial Genomics 10:001224.

12. Uruén C, Fernandez A, Arnal JL, del Pozo M, Amoribieta MC, de Blas I, Jurado P, Calvo JH, Gottschalk M, González-Vázquez LD, Arenas M, Marín CM, Arenas J. 2024. Genomic and phenotypic analysis of invasive Streptococcus suis isolated in Spain reveals genetic diversification and associated virulence traits. Veterinary Research 55:11.

13. Li K, Lacouture S, Lewandowski E, Thibault E, Gantelet H, Gottschalk M, Fittipaldi N. 2024. Molecular characterization of Streptococcus suis isolates recovered from diseased pigs in Europe. Vet Res 55:117.

14. Clokie MRJ, Millard AD, Letarov AV, Heaphy S. 2011. Phages in nature. Bacteriophage 10.4161/bact.1.1.14942.

15. Thanki AM, Hooton S, Whenham N, Salter MG, Bedford MR, O’Neill HVM, Clokie MRJ. 2023. A bacteriophage cocktail delivered in feed significantly reduced Salmonella colonization in challenged broiler chickens. Emerging Microbes & Infections.

16. Thanki AM, Osei EK, Whenham N, Salter MG, Bedford MR, Masey O’Neill HV, Clokie MRJ. 2024. Broad host range phages target global Clostridium perfringens bacterial strains and clear infection in five-strain model systems. Microbiology Spectrum 12:e03784–23.

17. Nick JA, Dedrick RM, Gray AL, Vladar EK, Smith BE, Freeman KG, Malcolm KC, Epperson LE, Hasan NA, Hendrix J, Callahan K, Walton K, Vestal B, Wheeler E, Rysavy NM, Poch K, Caceres S, Lovell VK, Hisert KB, Moura VC de, Chatterjee D, De P, Weakly N, Martiniano SL, Lynch DA, Daley CL, Strong M, Jia F, Hatfull GF, Davidson RM. 2022. Host and pathogen response to bacteriophage engineered against Mycobacterium abscessus lung infection. Cell 185:1860–1874.e12.

18. Pirnay J-P, Djebara S, Steurs G, Griselain J, Cochez C, De Soir S, Glonti T, Spiessens A, Vanden Berghe E, Green S, Wagemans J, Lood C, Schrevens E, Chanishvili N, Kutateladze M, de Jode M, Ceyssens P-J, Draye J-P, Verbeken G, De Vos D, Rose T, Onsea J, Van Nieuwenhuyse B, Soentjens P, Lavigne R, Merabishvili M. 2024. Personalized bacteriophage therapy outcomes for 100 consecutive cases: a multicentre, multinational, retrospective observational study. Nat Microbiol 9:1434–1453.

19. Ma YL, Lu CP. 2008. Isolation and identification of a bacteriophage capable of infecting *Streptococcus suis* type 2 strains. Veterinary Microbiology 132:340– 347.

20. Osei EK, Mahony J, Kenny JG. 2022. From Farm to Fork: Streptococcus suis as a Model for the Development of Novel Phage-Based Biocontrol Agents. 9. Viruses 14:1996.

21. Arroyo-Moreno S, Cummings M, Corcoran DB, Coffey A, McCarthy RR. 2022. Identification and characterization of novel endolysins targeting Gardnerella vaginalis biofilms to treat bacterial vaginosis. npj Biofilms Microbiomes 8:1–12.

22. Mondal SI, Akter A, Draper LA, Ross RP, Hill C. 2021. Characterization of an Endolysin Targeting Clostridioides difficile That Affects Spore Outgrowth. International Journal of Molecular Sciences 22:5690.

23. Elst NV, Linden SB, Lavigne R, Meyer E, Briers Y, Nelson DC. 2020. Characterization of the Bacteriophage-Derived Endolysins PlySs2 and PlySs9 with In Vitro Lytic Activity against Bovine Mastitis Streptococcus uberis. Antibiotics 9:621.

24. Vázquez R, Domenech M, Iglesias-Bexiga M, Menéndez M, García P. 2017. Csl2, a novel chimeric bacteriophage lysin to fight infections caused by Streptococcus suis, an emerging zoonotic pathogen. Sci Rep 7:16506.

25. Atack JM, Weinert LA, Tucker AW, Husna AU, Wileman TM, F. Hadjirin N, Hoa NT, Parkhill J, Maskell DJ, Blackall PJ, Jennings MP. 2018. Streptococcus suis contains multiple phase-variable methyltransferases that show a discrete lineage distribution. Nucleic Acids Research 46:11466–11476.

26. Okura M, Lachance C, Osaki M, Sekizaki T, Maruyama F, Nozawa T, Nakagawa I, Hamada S, Rossignol C, Gottschalk M, Takamatsu D. 2020. Development of a Two-Step Multiplex PCR Assay for Typing of Capsular Polysaccharide Synthesis Gene Clusters of Streptococcus suis. Journal of Clinical Microbiology 52:1714–1719.

27. Athey TBT, Teatero S, Lacouture S, Takamatsu D, Gottschalk M, Fittipaldi N. 2016. Determining Streptococcus suis serotype from short-read whole-genome sequencing data. BMC Microbiology 16:162.

28. 2018. Open-access bacterial population genomics: BIGSdb software, the … Wellcome Open Research | Open Access Publishing Platform. https://wellcomeopenresearch.org/articles/3-124/v1?s3BucketUrl=https%3A%2F%2Fwellcomeopenresearch.s3.eu-west-1.amazonaws.com&gtmKey=GTM-5P673KJ&submissionUrl=%2Ffor-authors%2Fpublish-your-research&otid=23226e40-fdd0-4acd-97a3-d9bad93befed&immUserUrl=https%3A%2F%2Fwor-proxy.f1krdev.com%2Feditor%2Fmember%2Fshow%2F. Retrieved 9 November 2024.

29. da Silva RT, de Souza Grilo MM, Magnani M, de Souza Pedrosa GT. 2021. Double-Layer Plaque Assay Technique for Enumeration of Virus Surrogates, p. 157–162. *In* Magnani, M (ed.), Detection and Enumeration of Bacteria, Yeast, Viruses, and Protozoan in Foods and Freshwater. Springer US, New York, NY.

30. Camargo AP, Roux S, Schulz F, Babinski M, Xu Y, Hu B, Chain PSG, Nayfach S, Kyrpides NC. 2024. Identification of mobile genetic elements with geNomad. Nat Biotechnol 42:1303–1312.

31. Bouras G, Nepal R, Houtak G, Psaltis AJ, Wormald P-J, Vreugde S. 2023. Pharokka: a fast scalable bacteriophage annotation tool. Bioinformatics 39:btac776.

32. Seemann T. 2024. tseemann/shovill. Perl.

33. CheckV assesses the quality and completeness of metagenome-assembled viral genomes | Nature Biotechnology. https://www.nature.com/articles/s41587-020-00774-7. Retrieved 9 November 2024.

34. Aziz RK, Bartels D, Best AA, DeJongh M, Disz T, Edwards RA, Formsma K, Gerdes S, Glass EM, Kubal M, Meyer F, Olsen GJ, Olson R, Osterman AL, Overbeek RA, McNeil LK, Paarmann D, Paczian T, Parrello B, Pusch GD, Reich C, Stevens R, Vassieva O, Vonstein V, Wilke A, Zagnitko O. 2008. The RAST Server: Rapid Annotations using Subsystems Technology. BMC Genomics 9:75.

35. Bouras G. 2024. gbouras13/phold. Python.

36. Shang J, Peng C, Liao H, Tang X, Sun Y. 2023. PhaBOX: a web server for identifying and characterizing phage contigs in metagenomic data. Bioinformatics Advances 3:vbad101.

37. VIBRANT: automated recovery, annotation and curation of microbial viruses, and evaluation of viral community function from genomic sequences | Microbiome | Full Text. https://microbiomejournal.biomedcentral.com/articles/10.1186/s40168-020-00867-0. Retrieved 9 November 2024.

38. DRAM for distilling microbial metabolism to automate the curation of microbiome function | Nucleic Acids Research | Oxford Academic. https://academic.oup.com/nar/article/48/16/8883/5884738. Retrieved 9 November 2024.

39. Tesson F, Hervé A, Mordret E, Touchon M, d’Humières C, Cury J, Bernheim A. 2022. Systematic and quantitative view of the antiviral arsenal of prokaryotes. Nat Commun 13:2561.

40. Sievers F, Higgins DG. 2017. Clustal Omega for making accurate alignments of many protein sequences. Protein Science : A Publication of the Protein Society 27:135.

41. vragh. 2022. vragh/seqvisr: v0.2.7 (v0.2.7). Zenodo.

42. ViPTree: the viral proteomic tree server | Bioinformatics | Oxford Academic. https://academic.oup.com/bioinformatics/article/33/15/2379/3096426?login=false. Retrieved 9 November 2024.

43. Millard A, Denise R, Lestido M, Thomas M, Turner D, Turner D, Sicheritz-Pontén T. 2024. taxmyPHAGE: Automated taxonomy of dsDNA phage genomes at the genus and species level. bioRxiv 10.1101/2024.08.09.606593.

44. VirClust—A Tool for Hierarchical Clustering, Core Protein Detection and Annotation of (Prokaryotic) Viruses. https://www.mdpi.com/1999-4915/15/4/1007. Retrieved 9 November 2024.

45. Kropinski AM. 2018. Practical Advice on the One-Step Growth Curve, p. 41–47. In Clokie, MRJ, Kropinski, AM, Lavigne, R (eds.), Bacteriophages: Methods and Protocols, Volume 3. Springer, New York, NY.

46. Jumper J, Evans R, Pritzel A, Green T, Figurnov M, Ronneberger O, Tunyasuvunakool K, Bates R, Žídek A, Potapenko A, Bridgland A, Meyer C, Kohl SAA, Ballard AJ, Cowie A, Romera-Paredes B, Nikolov S, Jain R, Adler J, Back T, Petersen S, Reiman D, Clancy E, Zielinski M, Steinegger M, Pacholska M, Berghammer T, Bodenstein S, Silver D, Vinyals O, Senior AW, Kavukcuoglu K, Kohli P, Hassabis D. 2021. Highly accurate protein structure prediction with AlphaFold. Nature 596:583–589.

47. Abramson J, Adler J, Dunger J, Evans R, Green T, Pritzel A, Ronneberger O, Willmore L, Ballard AJ, Bambrick J, Bodenstein SW, Evans DA, Hung C-C, O’Neill M, Reiman D, Tunyasuvunakool K, Wu Z, Žemgulytė A, Arvaniti E, Beattie C, Bertolli O, Bridgland A, Cherepanov A, Congreve M, Cowen-Rivers AI, Cowie A, Figurnov M, Fuchs FB, Gladman H, Jain R, Khan YA, Low CMR, Perlin K, Potapenko A, Savy P, Singh S, Stecula A, Thillaisundaram A, Tong C, Yakneen S, Zhong ED, Zielinski M, Žídek A, Bapst V, Kohli P, Jaderberg M, Hassabis D, Jumper JM. 2024. Accurate structure prediction of biomolecular interactions with AlphaFold 3. Nature 630:493–500.

48. Emsley P, Lohkamp B, Scott WG, Cowtan K. 2010. Features and development of Coot. Acta Cryst D 66:486–501.

49. van Kempen M, Kim SS, Tumescheit C, Mirdita M, Lee J, Gilchrist CLM, Söding J, Steinegger M. 2024. Fast and accurate protein structure search with Foldseek. Nat Biotechnol 42:243–246.

50. Corpet F. 1988. Multiple sequence alignment with hierarchical clustering. Nucleic Acids Res 16:10881–10890.

51. Pettersen EF, Goddard TD, Huang CC, Couch GS, Greenblatt DM, Meng EC, Ferrin TE. 2004. UCSF Chimera—A visualization system for exploratory research and analysis. Journal of Computational Chemistry 25:1605–1612.

52. Meng X, Shi Y, Ji W, Meng X, Zhang J, Wang H, Lu C, Sun J, Yan Y. 2011. Application of a Bacteriophage Lysin To Disrupt Biofilms Formed by the Animal Pathogen Streptococcus suis. Applied and Environmental Microbiology 77:8272–8279.

53. Ji W, Huang Q, Sun L, Wang H, Yan Y, Sun J. 2015. A novel endolysin disrupts Streptococcus suis with high efficiency. FEMS Microbiology Letters 362:fnv205.

54. Combined Antibacterial Activity of Phage Lytic Proteins Holin and Lysin from Streptococcus suis Bacteriophage SMP | Current Microbiology. https://link.springer.com/article/10.1007/s00284-012-0119-2. Retrieved 28 November 2024.

55. Goulet A, Joos R, Lavelle K, Van Sinderen D, Mahony J, Cambillau C. 2022. A structural discovery journey of streptococcal phages adhesion devices by AlphaFold2. Front Mol Biosci 9.

56. Goulet A, Spinelli S, Mahony J, Cambillau C. 2020. Conserved and Diverse Traits of Adhesion Devices from Siphoviridae Recognizing Proteinaceous or Saccharidic Receptors. 5. Viruses 12:512.

57. Lavelle K, Goulet A, McDonnell B, Spinelli S, van Sinderen D, Mahony J, Cambillau C. 2020. Revisiting the host adhesion determinants of Streptococcus thermophilus siphophages. Microbial Biotechnology 13:1765– 1779.

58. Veesler D, Robin G, Lichière J, Auzat I, Tavares P, Bron P, Campanacci V, Cambillau C. 2010. Crystal Structure of Bacteriophage SPP1 Distal Tail Protein (gp19.1): A BASEPLATE HUB PARADIGM IN GRAM-POSITIVE INFECTING PHAGES *. Journal of Biological Chemistry 285:36666–36673.

59. Veesler D, Cambillau C. 2011. A Common Evolutionary Origin for Tailed-Bacteriophage Functional Modules and Bacterial Machineries. Microbiology and Molecular Biology Reviews 75:423–433.

60. Flayhan A, Vellieux FMD, Lurz R, Maury O, Contreras-Martel C, Girard E, Boulanger P, Breyton C. 2014. Crystal Structure of pb9, the Distal Tail Protein of Bacteriophage T5: a Conserved Structural Motif among All Siphophages. Journal of Virology 88:820–828.

61. Dieterle ME, Fina Martin J, Durán R, Nemirovsky SI, Sanchez Rivas C, Bowman C, Russell D, Hatfull GF, Cambillau C, Piuri M. 2016. Characterization of prophages containing “evolved” Dit/Tal modules in the genome of Lactobacillus casei BL23. Appl Microbiol Biotechnol 100:9201–9215.

62. Dieterle M-E, Spinelli S, Sadovskaya I, Piuri M, Cambillau C. 2017. Evolved distal tail carbohydrate binding modules of actobacillus phage J-1: a novel type of anti-receptor widespread among lactic acid bacteria phages. Molecular Microbiology 104:608–620.

63. Sciara G, Bebeacua C, Bron P, Tremblay D, Ortiz-Lombardia M, Lichière J, van Heel M, Campanacci V, Moineau S, Cambillau C. 2010. Structure of lactococcal phage p2 baseplate and its mechanism of activation. Proceedings of the National Academy of Sciences 107:6852–6857.

64. Mahony J, Goulet A, van Sinderen D, Cambillau C. 2023. Partial Atomic Model of the Tailed Lactococcal Phage TP901-1 as Predicted by AlphaFold2: Revelations and Limitations. 12. Viruses 15:2440.

65. Stockdale SR, Mahony J, Courtin P, Chapot-Chartier M-P, van Pijkeren J-P, Britton RA, Neve H, Heller KJ, Aideh B, Vogensen FK, van Sinderen D. 2013. The Lactococcal Phages Tuc2009 and TP901-1 Incorporate Two Alternate Forms of Their Tail Fiber into Their Virions for Infection Specialization* [S]. Journal of Biological Chemistry 288:5581–5590.

66. São-José C, Lhuillier S, Lurz R, Melki R, Lepault J, Santos MA, Tavares P. 2006. The Ectodomain of the Viral Receptor YueB Forms a Fiber That Triggers Ejection of Bacteriophage SPP1 DNA*. Journal of Biological Chemistry 281:11464–11470.

67. Spinelli S, Campanacci V, Blangy S, Moineau S, Tegoni M, Cambillau C. 2006. Modular Structure of the Receptor Binding Proteins of *Lactococcus lactis* Phages. Journal of Biological Chemistry 281:14256–14262.

68. Spinelli S, Desmyter A, Verrips CT, de Haard HJW, Moineau S, Cambillau C. 2006. Lactococcal bacteriophage p2 receptor-binding protein structure suggests a common ancestor gene with bacterial and mammalian viruses. Nat Struct Mol Biol 13:85–89.

69. Dunne M, Hupfeld M, Klumpp J, Loessner MJ. 2018. Molecular Basis of Bacterial Host Interactions by Gram-Positive Targeting Bacteriophages. 8. Viruses 10:397.

70. Caulton SG, Lambert C, Tyson J, Radford P, Al-Bayati A, Greenwood S, Banks EJ, Clark C, Till R, Pires E, Sockett RE, Lovering AL. 2024. Bdellovibrio bacteriovorus uses chimeric fibre proteins to recognize and invade a broad range of bacterial hosts. Nat Microbiol 9:214–227.

71. Andres D, Gohlke U, Broeker NK, Schulze S, Rabsch W, Heinemann U, Barbirz S, Seckler R. 2013. An essential serotype recognition pocket on phage P22 tailspike protein forces Salmonella enterica serovar Paratyphi A O-antigen fragments to bind as nonsolution conformers. Glycobiology 23:486–494.

72. Mavrich TN, Casey E, Oliveira J, Bottacini F, James K, Franz CMAP, Lugli GA, Neve H, Ventura M, Hatfull GF, Mahony J, van Sinderen D. 2018. Characterization and induction of prophages in human gut-associated Bifidobacterium hosts. Sci Rep 8:12772.

73. Deecker SR, Urbanus ML, Nicholson B, Ensminger AW. Legionella pneumophila CRISPR-Cas Suggests Recurrent Encounters with One or More Phages in the Family Microviridae. Appl Environ Microbiol 87:e00467–21.

74. Nale JY, Spencer J, Hargreaves KR, Buckley AM, Trzepiński P, Douce GR, Clokie MRJ. 2016. Bacteriophage Combinations Significantly Reduce Clostridium difficile Growth In Vitro and Proliferation In Vivo. Antimicrob Agents Chemother 60:968–981.

75. Tram G, Jen FE-C, Phillips ZN, Timms J, Husna A-U, Jennings MP, Blackall PJ, Atack JM. 2021. Streptococcus suis Encodes Multiple Allelic Variants of a Phase-Variable Type III DNA Methyltransferase, ModS, That Control Distinct Phasevarions. mSphere 6:10.1128/msphere.00069-21.

76. Roodsant TJ, van der Putten B, Brizuela J, Coolen JPM, Baltussen TJH, Schipper K, Pannekoek Y, van der Ark KCH, Schultsz C. The streptococcal phase-variable type I restriction modification system SsuCC20p dictates the methylome of Streptococcus suis impacting the transcriptome and virulence in a zebrafish larvae infection model. mBio 15:e02259–23.

77. Cayrou C, Barratt NA, Ketley JM, Bayliss CD. 2021. Phase Variation During Host Colonization and Invasion by Campylobacter jejuni and Other Campylobacter Species. Front Microbiol 12.

78. Sekulovic O, Bedoya MO, Fivian-Hughes AS, Fairweather NF, Fortier L-C. 2015. The C lostridium difficile cell wall protein CwpV confers phase-variable phage resistance. Molecular Microbiology 98:329.

79. Turkington CJR, Morozov A, Clokie MRJ, Bayliss CD. 2019. Phage-Resistant Phase-Variant Sub-populations Mediate Herd Immunity Against Bacteriophage Invasion of Bacterial Meta-Populations. Front Microbiol 10.

80. Tang F, Bossers A, Harders F, Lu C, Smith H. 2013. Comparative genomic analysis of twelve *Streptococcus suis* (pro)phages. Genomics 101:336–344.

81. Zeng Z, Liu X, Yao J, Guo Y, Li B, Li Y, Jiao N, Wang X. 2016. Cold adaptation regulated by cryptic prophage excision in Shewanella oneidensis. ISME J 10:2787–2800.

82. Prophage Induction by High Temperature in Thermosensitive dna Mutants Lysogenic for Bacteriophage Lambda. https://journals.asm.org/doi/epdf/10.1128/jvi.11.6.879-885.1973. Retrieved 25 November 2024.

83. Spontaneously induced prophages are abundant in a naturally evolved bacterial starter culture and deliver competitive advantage to the host | BMC Microbiology | Full Text. https://bmcmicrobiol.biomedcentral.com/articles/10.1186/s12866-018-1229-1. Retrieved 25 November 2024.

84. Molecular Ecology | Molecular Genetics Journal | Wiley Online Library. https://onlinelibrary.wiley.com/doi/full/10.1111/mec.16638. Retrieved 25 November 2024.

85. Gascón I, Lázaro JM, Salas M. 2000. Differential functional behavior of viral ϕ29, Nf and GA-1 SSB proteins. Nucleic Acids Research 28:2034–2042.

86. A broadly distributed predicted helicase/nuclease confers phage resistance via abortive infection - ScienceDirect. https://www.sciencedirect.com/science/article/pii/S1931312823000355. Retrieved 25 November 2024.

87. Sasaki T, Takita S, Fujishiro T, Shintani Y, Nojiri S, Yasui R, Yonesaki T, Otsuka Y. 2023. Phage single-stranded DNA-binding protein or host DNA damage triggers the activation of the AbpAB phage defense system. mSphere 8:e00372–23.

88. Baker ML, Jiang W, Rixon FJ, Chiu W. 2005. Common Ancestry of Herpesviruses and Tailed DNA Bacteriophages. J Virol 79:14967–14970.

89. Kizziah JL, Manning KA, Dearborn AD, Dokland T. 2020. Structure of the host cell recognition and penetration machinery of a Staphylococcus aureus bacteriophage. PLoS Pathog 16:e1008314.

90. Harel J, Martinez G, Nassar A, Dezfulian H, Labrie SJ, Brousseau R, Moineau S, Gottschalk M. 2003. Identification of an Inducible Bacteriophage in a Virulent Strain of Streptococcus suis Serotype 2. Infection and Immunity 71:6104–6108.

91. Drivers and consequences of bacteriophage host range | FEMS Microbiology Reviews | Oxford Academic. https://academic.oup.com/femsre/article/47/4/fuad038/7221647. Retrieved 25 November 2024.

92. Coevolutionary diversification creates nested-modular structure in phage– bacteria interaction networks - PMC. https://pmc.ncbi.nlm.nih.gov/articles/PMC3915849/. Retrieved 25 November 2024.

93. Gurney J, Aldakak L, Betts A, Gougat-Barbera C, Poisot T, Kaltz O, Hochberg ME. 2017. Network structure and local adaptation in co-evolving bacteria– phage interactions. Molecular Ecology 26:1764–1777.

94. Bull JJ, Wichman HA, Krone SM, Molineux IJ. 2024. Controlling Recombination to Evolve Bacteriophages. Cells 13:585.

95. Peters TL, Song Y, Bryan DW, Hudson LK, Denes TG. 2020. Mutant and Recombinant Phages Selected from In Vitro Coevolution Conditions Overcome Phage-Resistant Listeria monocytogenes. Appl Environ Microbiol 86:e02138–20.

96. Abedon ST, LeJeune JT. 2007. Why Bacteriophage Encode Exotoxins and other Virulence Factors. Evol Bioinform Online 1:97–110.

97. Capparelli R, Nocerino N, Lanzetta R, Silipo A, Amoresano A, Giangrande C, Becker K, Blaiotta G, Evidente A, Cimmino A, Iannaccone M, Parlato M, Medaglia C, Roperto S, Roperto F, Ramunno L, Iannelli D. 2010. Bacteriophage-Resistant Staphylococcus aureus Mutant Confers Broad Immunity against Staphylococcal Infection in Mice. PLOS ONE 5:e11720.

98. Fujiki J, Nakamura K, Nakamura T, Iwano H. 2023. Fitness Trade-Offs between Phage and Antibiotic Sensitivity in Phage-Resistant Variants: Molecular Action and Insights into Clinical Applications for Phage Therapy. 21. International Journal of Molecular Sciences 24:15628.

99. Collateral sensitivity increases the efficacy of a rationally designed bacteriophage combination to control Salmonella enterica | Journal of Virology. https://journals.asm.org/doi/10.1128/jvi.01476-23. Retrieved 25 November 2024.

100. Yoo S, Lee K-M, Kim N, Vu TN, Abadie R, Yong D. 2023. Designing phage cocktails to combat the emergence of bacteriophage-resistant mutants in multidrug-resistant Klebsiella pneumoniae. Microbiology Spectrum 12:e01258–23.

101. Niu YD, Liu H, Du H, Meng R, Sayed Mahmoud E, Wang G, McAllister TA, Stanford K. 2021. Efficacy of Individual Bacteriophages Does Not Predict Efficacy of Bacteriophage Cocktails for Control of Escherichia coli O157. Front Microbiol 12.

102. Thanki AM, Mignard G, Atterbury RJ, Barrow P, Millard AD, Clokie MRJ. 2022. Prophylactic Delivery of a Bacteriophage Cocktail in Feed Significantly Reduces Salmonella Colonization in Pigs. Microbiology Spectrum 10:e00422–22.

103. Zhang Y, Chu M, Liao Y-T, Salvador A, Wu VCH. 2024. Characterization of two novel Salmonella phages having biocontrol potential against Salmonella spp. in gastrointestinal conditions. Sci Rep 14:12294.

