## Supplementary material for "Isolation of phages infecting the zoonotic pathogen *Streptococcus suis* reveals novel structural and genomic characteristics": Table S1-S3 and Fig S1-S3

**Table S1: Strains used in this study**

| **Strain** | **Host range** | | **Serotype** | **ST** | **Year** | **Pathotype** | **Country** | **Accession No.** |
| --- | --- | --- | --- | --- | --- | --- | --- | --- |
|  | **Bonnie** | **Clyde** |  |  |  |  |  |  |
| 94_8576_4887* | - | + | 14 | 28 | 1994 | Respiratory | Canada | PRJNA628943 |
| 89_5046* | - | + | 2 | 25 | 1989 | Systemic | Canada | PRJNA628943 |
| 89_6891_2* | - | + | 2 | 25 | 1989 | Respiratory | Canada | PRJNA628943 |
| 90_2741_7 | - | + | 2 | 25 | 1990 | Systemic | Canada | PRJNA628943 |
| 1602951 | - | + | 2 | 25 | 2014 | Respiratory | Canada | PRJNA628943 |
| 1596201* | - | + | 14 | 1 | 2014 | Systemic | Canada | PRJNA628943 |
| 1463646* | - | + | 14 | 1 | 2013 | Systemic | Canada | PRJNA628943 |
| 1667796 | - | + | 2 | NF | 2014 | Systemic | Canada | PRJNA628943 |
| 1093404* | - | + | 27 | NF | 2008 | Systemic | Canada | PRJNA628943 |
| 1162351* | - | - | 24 | NF | 2009 | Respiratory | Canada | PRJNA628943 |
| 1269330* | - | - | 21 | NF | 2011 | Systemic | Canada | PRJNA628943 |
| 21137_DNR4 | - | + | 2 | 28 | 2018 | Respiratory | Denmark | PRJNA1009400 |
| 21136_DNR3 | - | + | 2 | 28 | 2018 | Respiratory | Denmark | PRJNA1009400 |
| 21135_DNR2* | - | + | 2 | 28 | 2018 | Respiratory | Denmark | PRJNA1009400 |
| 21169_DNR36*↑ | + | + | 2 | 28 | 2018 | Respiratory | Denmark | PRJNA1009400 |
| 21171_DNR38*↑ | + | + | 2 | 1 | 2018 | Respiratory | Denmark | PRJNA1009400 |
| 21172_DNR39 | + | + | 2 | 1 | 2018 | Respiratory | Denmark | PRJNA1009400 |
| 21173_DNR40 | - | + | 2 | 28 | 2018 | Respiratory | Denmark | PRJNA1009400 |
| 21176_DNR43 | - | + | 2 | 28 | 2018 | Respiratory | Denmark | PRJNA1009400 |
| 21225_DNS42 | - | - | 7 | 29 | 2018 | Systemic | Denmark | PRJNA1009400 |
| 21224_DNS41 | - | - | 2 | 28 | 2018 | Systemic | Denmark | PRJNA1009400 |
| 21223_DNS40 | - | - | 7 | 29 | 2018 | Systemic | Denmark | PRJNA1009400 |
| 21222_DNS39 | - | - | 7 | 29 | 2018 | Systemic | Denmark | PRJNA1009400 |
| D1* | - | + | 2 | 1 | 2019 | Unknown | Ireland | N/A |
| D3* | + | + | 2 | 1 | 2019 | Respiratory | Ireland | N/A |
| D5* | - | + | 2 | 1 | 2019 | Systemic | Ireland | N/A |
| D6 * | + | + | 2 | 1 | 2019 | Systemic | Ireland | N/A |
| D7* | + | + | 2 | 1 | 2019 | Unknown | Ireland | N/A |
| D8*↓ | - | + | 2 | 28 | 2012 | Respiratory | Ireland | N/A |
| D10* | - | + | 2 | 1 | 2012 | Systemic | Ireland | N/A |
| D14* | - | + | 2 | 1 | 2011 | Systemic | Ireland | N/A |
| D16* | - | + | 14 | 1 | 2011 | Unknown | Ireland | N/A |
| D17 | + | + | 2 | 1 | 2011 | Systemic | Ireland | N/A |
| D19 | - | + | 2 | 1 | 2010 | Respiratory | Ireland | N/A |
| D23* | - | + | 14 | 1 | 2008 | Unknown | Ireland | N/A |
| D24 | - | + | 2 | 1 | 2012 | Respiratory | Ireland | N/A |
| D29 | - | + | 7 | 29 | 2008 | Respiratory | Ireland | N/A |
| D31* | - | + | 4 | 856 | 2008 | Unknown | Ireland | N/A |
| D32 | + | + | 2 | 2629 | 2007 | Unknown | Ireland | N/A |
| D35 | - | + | 2 | 2646 | 2019 | Respiratory | Ireland | N/A |
| D42* | - | + | 16 | 2632 | 2019 | Respiratory | Ireland | N/A |
| D46 | + | + | 2 | 1 | 2019 | Respiratory | Ireland | N/A |
| D49* | - | + | 16 | 2632 | 2019 | Systemic | Ireland | N/A |
| D52↓ | + | + | 2 | 1 | 2018 | Systemic | Ireland | N/A |
| D57 | - | + | 2 | 2646 | 2019 | Respiratory | Ireland | N/A |
| D58* | + | + | 14 | 1 | 2019 | Systemic | Ireland | N/A |
| D68 | + | + | 2 | 1 | 2020 | Unknown | Ireland | N/A |
| D71*↑ | + | + | 14 | 1 | 2020 | Systemic | Ireland | N/A |
| D75*↓ | - | + | 9 | 2640 | 2020 | Respiratory | Ireland | N/A |
| D78 | - | + | 2 | 1 | 2020 | Respiratory | Ireland | N/A |
| D79 | - | + | 2 | 25 | 2020 | Systemic | Ireland | N/A |
| D84 | - | + | 2 | 2641 | 2021 | Unknown | Ireland | N/A |
| D93 | - | + | 2 | 28 | 2021 | Systemic | Ireland | N/A |
| D94*↓ | - | + | 14 | 124 | 2021 | Systemic | Ireland | N/A |
| D2 | - | - | 7 | 29 | 2019 | Systemic | Ireland | N/A |
| D4* | - | - | 9 | 2627 | 2019 | Systemic | Ireland | N/A |
| D9 | - | - | 2 | 28 | 2012 | Respiratory | Ireland | N/A |
| D11 | - | - | 7 | 29 | 2010 | Respiratory | Ireland | N/A |
| D12* | - | - | 9 | 2630 | 2010 | Respiratory | Ireland | N/A |
| D13* | - | - | 10 | 2626 | 2009 | Systemic | Ireland | N/A |
| D21* | - | - | 7 | 29 | 2010 | Systemic | Ireland | N/A |
| D22* | - | - | 9 | 2627 | 2009 | Systemic | Ireland | N/A |
| D25* | - | - | 9 | 2630 | 2021 | Unknown | Ireland | N/A |
| D26 | - | - | 2 | 28 | 2008 | Respiratory | Ireland | N/A |
| D27 | - | - | unknown | 2628 | 2007 | Unknown | Ireland | N/A |
| D28 | - | - | 2 | 2646 | 2005 | Respiratory | Ireland | N/A |
| D30 | - | - | 2 | 2646 | 2008 | Unknown | Ireland | N/A |
| D33* | - | - | 3 | 27 | 2010 | Respiratory | Ireland | N/A |
| D36* | - | - | 9 | 2627 | 2019 | Unknown | Ireland | N/A |
| D37 | - | - | 2 | 2649 | 2019 | Systemic | Ireland | N/A |
| D38* | - | - | 28 | 2631 | 2019 | Systemic | Ireland | N/A |
| D39* | - | - | 9 | 16 | 2019 | Respiratory | Ireland | N/A |
| D41* | - | - | 9 | 16 | 2019 | Systemic | Ireland | N/A |
| D100* | - | - | 9 | 2621 | 2021 | Systemic | Ireland | N/A |
| D101* | - | - | 3 | 94 | 2020 | Systemic | Ireland | N/A |
| D102 | - | - | 9 | 2624 | 2020 | Systemic | Ireland | N/A |
| D103 | - | - | 9 | 2630 | 2020 | Unknown | Ireland | N/A |
| D104* | - | - | 12 | 2633 | 2020 | Systemic | Ireland | N/A |
| D105* | - | - | 12 | 2634 | 2020 | Systemic | Ireland | N/A |
| D106 | - | - | unknown | 2635 | 2020 | Respiratory | Ireland | N/A |
| D107 | - | - | 9 | 2630 | 2021 | Unknown | Ireland | N/A |
| D108 | - | - | 9 | 2630 | 2021 | Unknown | Ireland | N/A |
| D109* | - | - | 11 | 2625 | 2022 | Respiratory | Ireland | N/A |
| D110* | - | - | 4 | 23 | 2022 | Respiratory | Ireland | N/A |
| 19858_M105281_R30* | - | + | 7 | 24 | 2017 | Respiratory | Spain | PRJNA1009400 |
| 19867_M106485_R39*↑↓ | + | + | 2 | 1 | 2018 | Respiratory | Spain | PRJNA1009400 |
| 19779_M101513_S1 | - | + | 2 | 1 | 2017 | Systemic | Spain | PRJNA1009400 |
| 19797_M104300_S19 | - | + | 2 | 1 | 2017 | Systemic | Spain | PRJNA1009400 |
| 19798_M104300_S20 | - | + | 2 | 1 | 2017 | Systemic | Spain | PRJNA1009400 |
| 19802_M105040_S24↑ | - | + | 2 | 1 | 2017 | Systemic | Spain | PRJNA1009400 |
| 19803_M105040_S25 | - | + | 2 | 1 | 2017 | Systemic | Spain | PRJNA1009400 |
| 19806_M105244_S28 | - | + | 2 | 1 | 2017 | Systemic | Spain | PRJNA1009400 |
| 19807_M105244_S29 | - | + | 2 | 1 | 2017 | Systemic | Spain | PRJNA1009400 |
| 19785_M102095_S7* | + | + | 2 | 1 | 2017 | Systemic | Spain | PRJNA1009400 |
| 19835_M103466_R7* | - | - | 23 | 17 | 2017 | Respiratory | Spain | PRJNA1009400 |
| 19852_M104890_R24 | - | - | 3 | 15 | 2017 | Respiratory | Spain | PRJNA1009400 |
| 19862_M106003_R34 | - | - | 3 | 15 | 2017 | Respiratory | Spain | PRJNA1009400 |
| 19874_M107244_R46 | - | - | 4 | 94 | 2018 | Respiratory | Spain | PRJNA1009400 |
| 19869_M106931_R41 | - | - | 9 | 123 | 2018 | Respiratory | Spain | PRJNA1009400 |
| 19871_M107169_R43* | - | - | 9 | 123 | 2018 | Respiratory | Spain | PRJNA1009400 |
| 4078  (*Streptococcus thermophilus*) | - | - | N/A | N/A | 1996 | N/A | Ireland | CP065477 |
| 90728  (*Streptococcus thermophilus*) | - | - | N/A | N/A | 1995 | N/A | Ireland | CP065479 |
| 3107  (*L. lactis cremoris*) | - | - | N/A | N/A | N/A | N/A | N/A | CP031538 |
| NZ9000  (*L. lactis cremoris*) | - | - | N/A | N/A | N/A | N/A | N/A | CP002094 |
| DH5α  (*E. coli*) | - | - | N/A | N/A | 2016 | N/A | France | CP025520 |

A total of 100 strains were tested used in host range tests. Susceptibility is scored as + (plaques/clear zone) or – (no plaques/clear zone). Abbreviations: NF, not found (sequence type undetermined). Pathotype: “Respiratory” refers to strains recovered from the respiratory tract of diseased pigs; “Systemic” refers to strains isolated from sites outside the respiratory tract; “Unknown” refers to strains of unspecified origin. *Strains (n=50) used in preliminary phage screening. ↑ indicates strains in multi-strain mix A, and ↓ indicates strains in multi-strain mix B. N/A indicates field is not applicable or the information is unknown for non-*S. suis* strains.

**Table S2: PCR primers and conditions for *S. suis* serotyping.**

| **Target *cps* group or type** | **Primer sequences (5′ to 3′)** |  | **Putative gene product(s) (HGb)** | **Size (bp) of products** |
| --- | --- | --- | --- | --- |
|  | **Forward** | **Reverse** |  |  |
| **For grouping PCR** |  |  |  |  |
| I | TGGTTCAAATATCAATGCTC | ATTGGTTGTGAGTGCATTG | Aminotransferase (HG41) | 933 |
| IId | TCAAAATACGCACCTAAGGC | CACTCACCTGCCCCAAGAC | *N*-acetylneuraminic acid synthase NeuB (HG10) | 823 |
| III*^e^*^,^*^f^* | TGATTTGGGTGAGACCATG | CTCATGCTGGATAACACGT | *N*-acetylfucosamine synthase FlnC (HG26) | 583 |
| IV | ACAGTCGGTCAAGATAATCG | TCAGCTTGGGTAATATCTGG | Initial sugar transferase (HG21) and aminotransferase (HG22) | 455 |
| Vg | GGAAAGATGGAGGACCAGC | CCAACCAGACTCATATCCCC | Initial sugar transferase (HG6) | 265 |
| III and VI*^e^*^,^*^h^* | GATGCCCCAAGCGATATGCC (F1), GACGCACCAAGTGATATGCC (F2) | GGACCAACAATGGCCATCTC (R1), GGTCCGACAATAGCCATTTC (R2) | Initial sugar transferase (HG8) | 146 |
| **For typing PCR** |  |  |  |  |
| *cps* group I |  |  |  |  |
| 3 | GGTTTTGATTGGTCTAGTTG | CTCTAAAGCTCGATATCTAC | Wzy polymerase (HG90) | 214 |
| 13 | TATGGTTAAAGGTGGAACTG | CCTTGTATATATTCCCTCCA | Wzy polymerase (HG148) | 408 |
| 18 | TAATGGGATAGTTGCGTTAC | ATACATAAAGTTGTCCTGCG | Wzy polymerase (HG170) | 617 |
| *cps* group II |  |  |  |  |
| 2 and 1/2 | TTAGCAACGTTGCCAATAAG | AATCCTCCATTAAAACCCTG | Wzy polymerase (HG54) | 173 |
| 6 | GCTCACTATTTTTACATTACAC | TATTACTCCGCCAAATACAG | Wzy polymerase (HG109) | 278 |
| 1 and 14 | TTAGACAGACACCTTATAGG | CTAGCTTCGTTACTTGATTC | Wzy polymerase (HG50) | 386 |
| 16 | AAGGTTATCCACGAAAGATG | TCCGGCAATATTCTTTCAAG | Wzy polymerase (HG362) | 494 |
| 27 | AGACACTGCTTGCATTATTG | TCAGAATTACTTCCTGTTGC | Wzy polymerase (HG246) | 655 |
| *cps* group III |  |  |  |  |
| 21 | TATCATATTGAGAATCTTCCC | TTGCGTAGCATACAAAGTTC | Wzy polymerase (HG194) | 160 |
| 28 | ATTATGTTGGTTGCAGAAGG | CGACTCAATTGTTGTAGTAG | Wzy polymerase (HG254) | 272 |
| 29 | TTCTGGGATTTTAGGAATGC | CATGAAATACGCACTTGTAC | Wzy polymerase (HG259) | 415 |
| 30 | TATTGCACTAGCTTCAGAAC | TGCATCCATAGTTGTATTCG | Wzy polymerase (HG264) | 568 |
| *cps* group IV |  |  |  |  |
| 4 | GACTATCTGTATACCCAAAC | TCCTTCCAAGTATTCTCTAG | Wzy polymerase (HG96) | 903 |
| 5 | ATCTTAGGAATGATTCGGAC | ACCAGATATCTGAGCAAATG | Wzy polymerase (HG103) | 720 |
| 7 | AACTACCTACCTGAACTTTG | AGTCTAAAAGTGATCGAGTC | Wzy polymerase (HG113) | 566 |
| 17 | TAGCATCAGTTTATACGAGG | TAGTTTATCTGTGACACACC | Wzy polymerase (HG164) | 455 |
| 19 | GTGTCGCAAATCAAGTATTG | AAGCTAGTACAACAAGCATG | Wzy polymerase (HG174) | 348 |
| 23 | TAATGTATGCTCTGTCACTG | AACGAAACGGAATAGTTTGC | Wzy polymerase (HG213) | 221 |
| *cps* group V |  |  |  |  |
| 8 | AAATAAGGTAGGAGCTACTC | ATCCAACCTTAGCTTTCTGT | Wzy polymerase (HG120) | 446 |
| 15 | ATCGTTTTGAGATTGAGTGG | TAAACGGATTCGGTTACTCA | Wzy polymerase (HG153) | 542 |
| 20 | TGTGGATTTCTGGGATAATC | TGTGGACGAATTACTACTTG | Wzy polymerase (HG182) | 698 |
| 22 | GCATTATCAGGATTCTTTCC | CCAATTGGGTGTTCAAAAAG | Wzy polymerase (HG200) | 296 |
| 25 | GTTTGCTCCGATCATAATAG | CCAGTAAAAGGACTCAATAC | Wzy polymerase (HG229) | 174 |
| *cps* group VI |  |  |  |  |
| 9 | GAAAGTAGGTATATCTCAGC | GGGCTATTAAAACTCCTATC | Wzy polymerase (HG123) | 368 |
| 10 | TTTCCCATTTGCTTATGGAC | GGAATAAAAACGATTGGGAG | Wzy polymerase (HG130) | 633 |
| 11 | ATGCGATTGCAACAATTGAC | AGGCATGAGTAATACATAGG | Wzy polymerase (HG138) | 833 |
| 12 | AACAGGTATTTCAGGATTGC | CTCGGATAAAGATAATCAGC | Wzy polymerase (HG140) | 131 |
| 24 | TACTGAGATTTATTGGGACG | AAGCGATTGGATTACATTGC | Wzy polymerase (HG220) | 224 |
| 26 | TTATACCGAAATTTTGTTGCC | CGTCAATCATATAAAGTGGG | Wzy polymerase (HG240) | 472 |
| 33 | GATGTTTTCAACAGGTGTAC | CAAAGTACCTATTTTCAGCG | Wzy polymerase (HG286) | 710 |
| *cps* group VII |  |  |  |  |
| 31 | ACAATCGTTTCTGCAATACG | GATGAAAACATCGTTGGTAG | Wzy polymerase (HG274) | 842 |
|  | ATCAGTAGTGGGAATAGTTG | TTTACTGTTTTTCGACCGTG | Initial sugar transferase (HG269) | 423 |
| 32 | AACCGCTGTTGAATTAAGAG | TTCGTTAGTTGAACTGTTCC | Wzy polymerase (HG281) | 570 |
|  | TAGGACTATGGTTCCTAATG | TATTCTAGTTCAAGTCGCTC | Initial sugar transferase (HG278) | 342 |
| 34 | AAGTTTCATTCGAGGACTTC | GTATATAACACCGCAAGAAG | Wzy polymerase (HG287) | 246 |
|  | ATACAGTGATGTCTTGCAAC | ATTGCTTTTTGACAATCGGC | Initial sugar transferase (HG290) | 701 |
| **For all PCR** |  |  |  |  |
| **Internal control** | GAGTTTGATCCTGGCTCAG | AGAAAGGAGGTGATCCAGCC |  | 1,542–1,553 |

Serotyping of S. suis strains was carried out as described by Okura et al (1). All PCR mixtures contained PCR Master Mix (2X) (ThermoFisher, cat# K0172). Each primer was added at a final concentration of 0.2μM. PCR conditions included an initial denaturation at 95°C for 3 minutes, followed by 35 cycles of denaturation at 95°C for 30 seconds, primmer annealing at 60°C (for grouping) or 58°C (for typing) for 90 seconds, and extension at 72°C for 90 seconds. Final extension was at 72°C for 10 minutes.

**Table S3: prophage PCR primers**

| Primer ID | Sequence 5’ to 3’ | Target (bp) |
| --- | --- | --- |
| TerF | ACGGCTATGCTATTCCACGG | Terminase  (988) |
| TerR | GCCGTGATGATTTCGCTGAC |  |
| 16SF | GAGTTTGATCCTGGCTCAG | 16s rRNA  (1542) |
| 16SR | AGAAAGGAGGTGATCCAGCC |  |
| *HolF | GCGTTGGTTGGATTTGTGTCT | Holin  (200) |
| *HolR | CTTTTGACCTGCTCCGCAAG |  |

Details of primers used for detection of prophage (Clyde). All PCR mixtures contained PCR Master Mix (2X) (ThermoFisher, cat# K0172). Each primer was added at a final concentration of 0.2μM. The following cycling conditions were used: initial denaturation at 95°C for 10 minutes, followed by 35 cycles of denaturation at 95°C for 30 seconds, annealing at 56°C for 30 seconds, and extension at 72°C for 90 seconds. Final extension was at 72°C for 10 minutes. Primers targeting Holin (*) are specific to Bonnie and used to distinguish between Bonnie and Clyde.

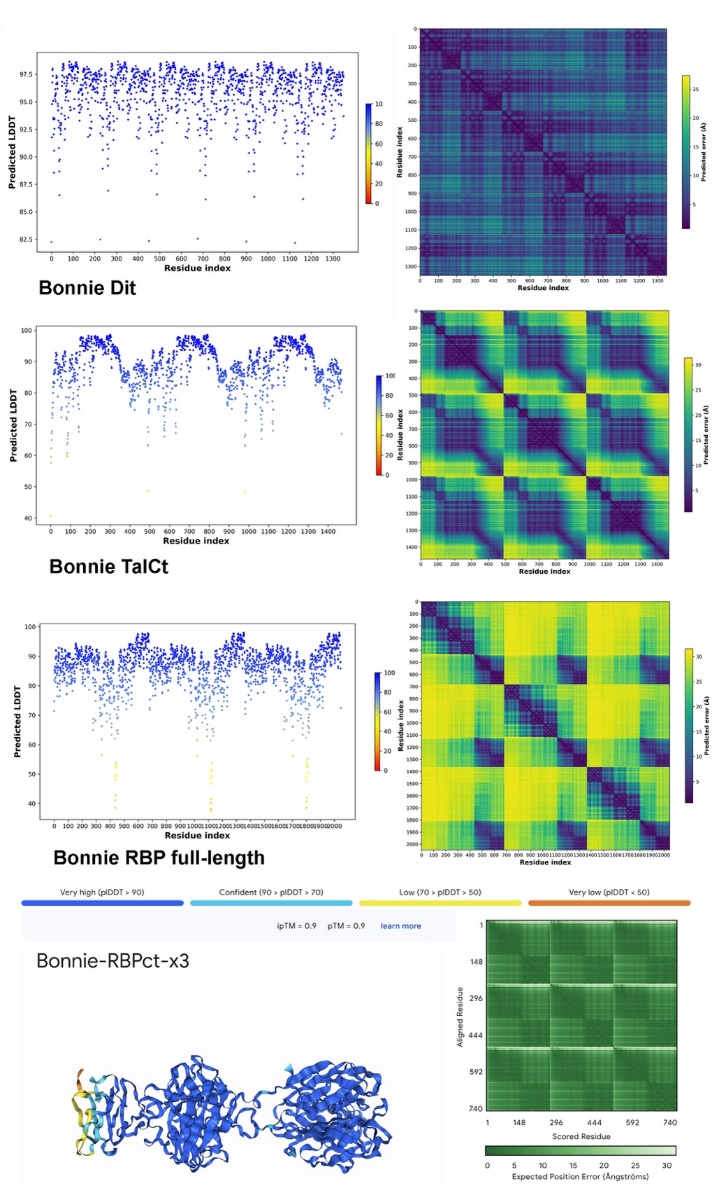

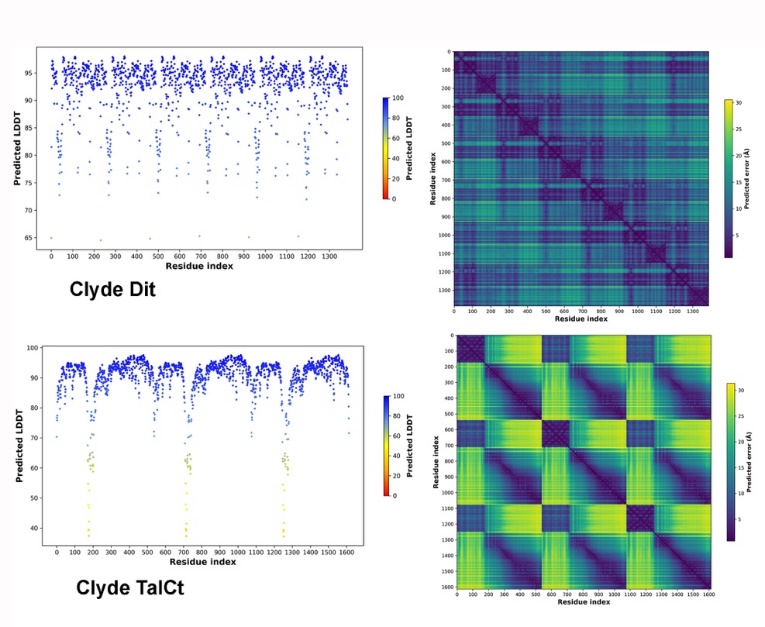

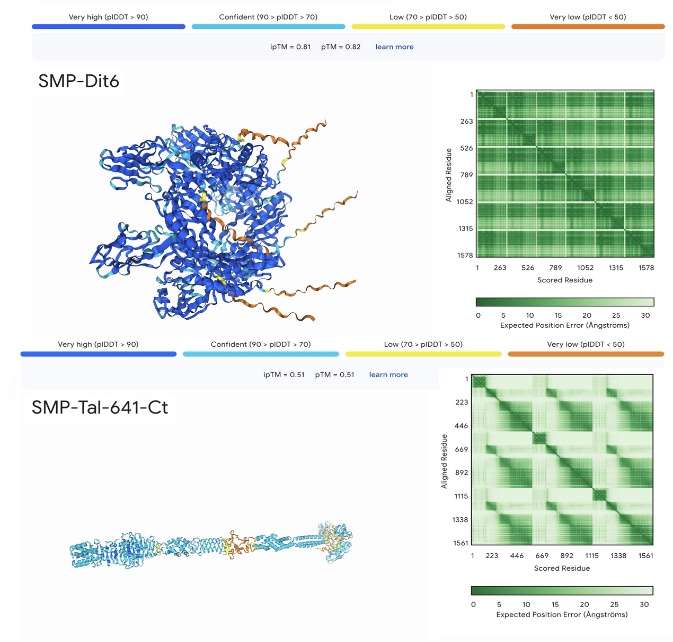

**B**

**A**

**C**

Fig. S1: Plots of the pLDDT and PEA statistics of **(A)** Bonnie, **(B)** Clyde, and **(C)** SMP

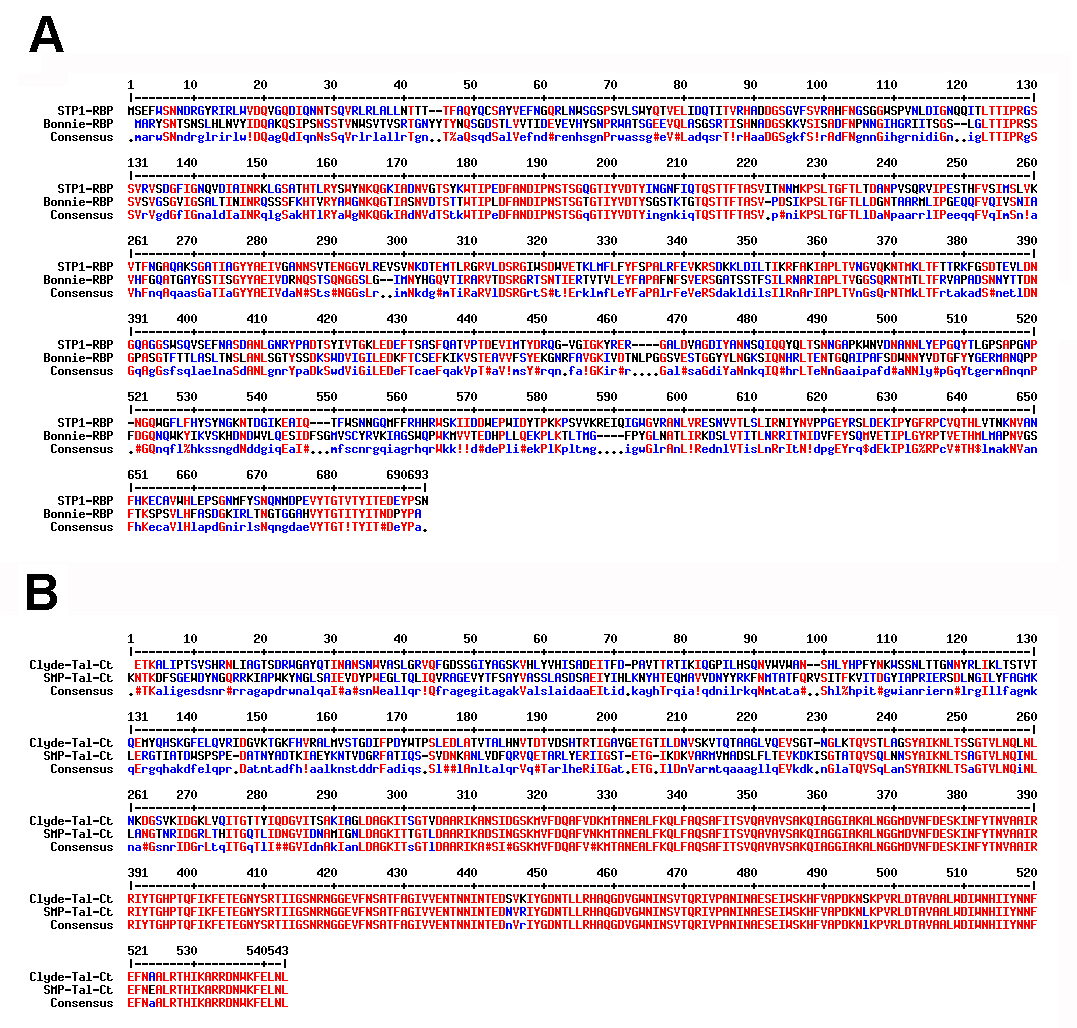

Fig. S2: Sequence alignments of the RBPs from phages STP1 and Bonnie **(A)** and of the Tal C-termini of phages Clyde and SMP **(B)**.

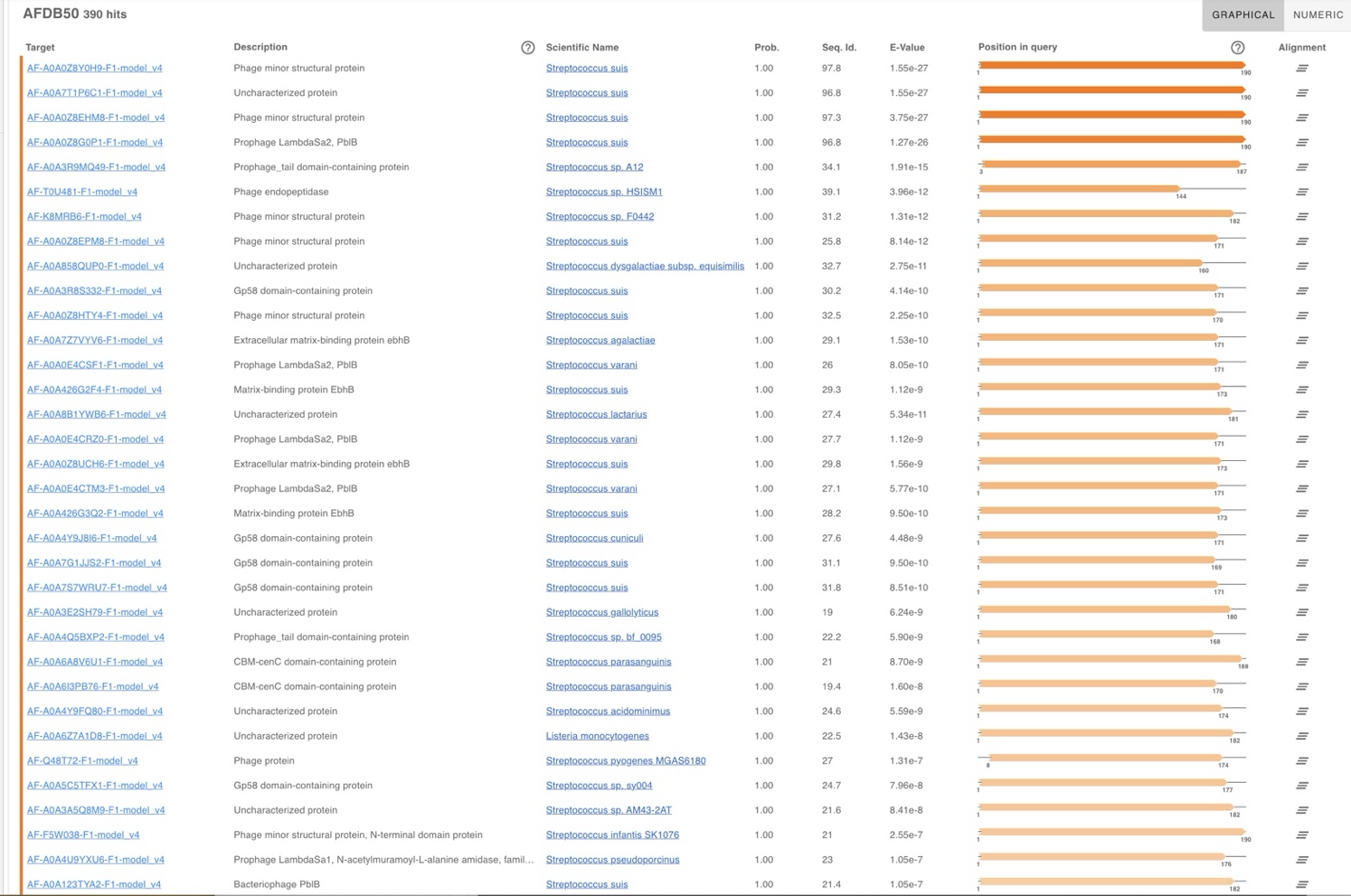

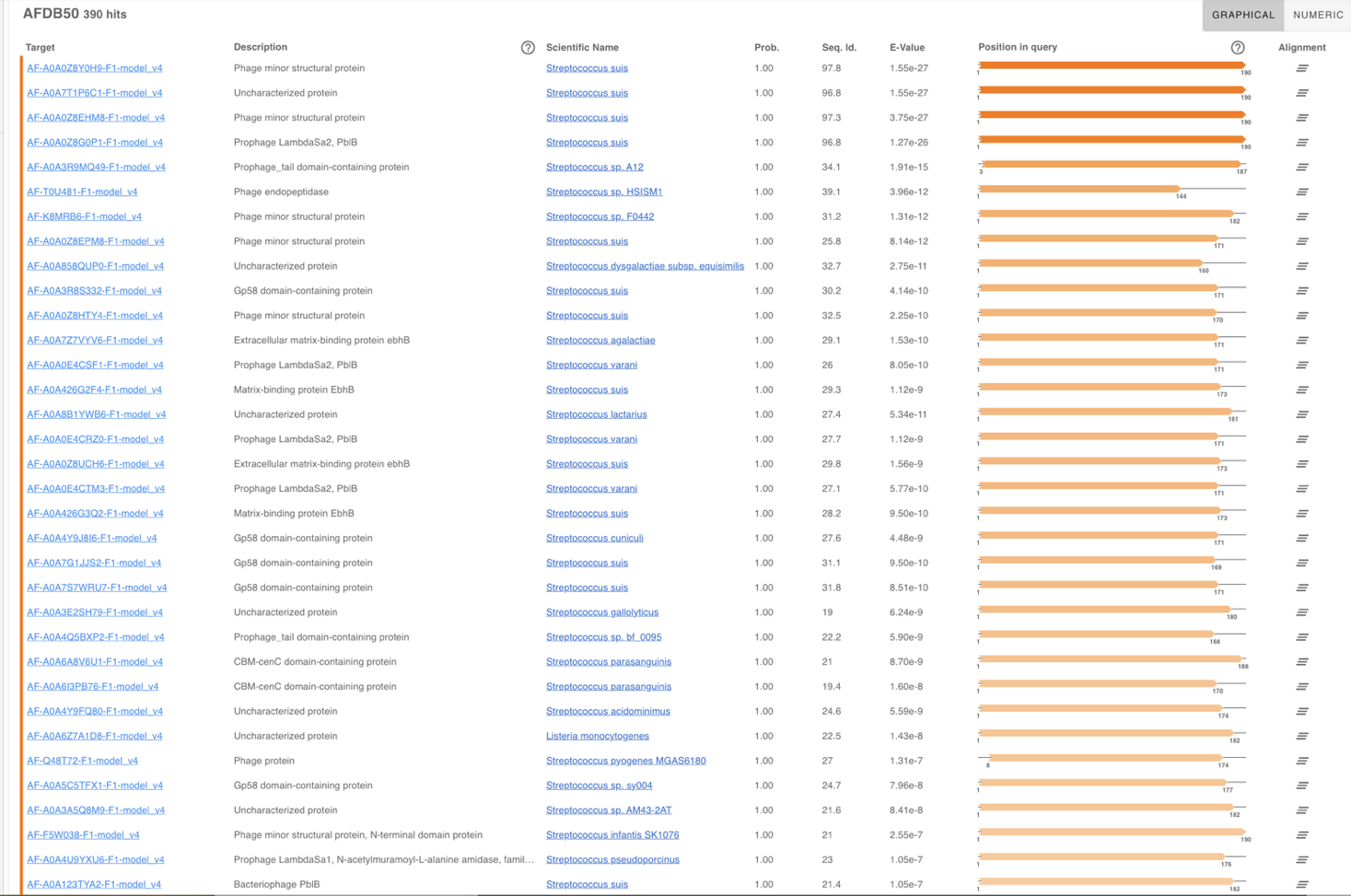

**A**

**B**

Fig. S3: Structural hits of **(A)** Clyde’s Tal C-terminus and **(B)** Bonnie’s RBP C-terminus in AlphaFold Database (AFDB)

References

1.	Okura M, Lachance C, Osaki M, Sekizaki T, Maruyama F, Nozawa T, Nakagawa I, Hamada S, Rossignol C, Gottschalk M, Takamatsu D. 2020. Development of a Two-Step Multiplex PCR Assay for Typing of Capsular Polysaccharide Synthesis Gene Clusters of Streptococcus suis. Journal of Clinical Microbiology 52:1714–1719.
